# Multiple origins of nematode-*Wolbachia* symbiosis in supergroup F and convergent loss of bacterioferritin in filarial *Wolbachia*

**DOI:** 10.1101/2021.03.23.436630

**Authors:** Amit Sinha, Zhiru Li, Catherine B. Poole, Laurence Ettwiller, Nathália F. Lima, Marcelo U. Ferreira, Fanny F. Fombad, Samuel Wanji, Clotilde K. S. Carlow

## Abstract

The intracellular endosymbiotic proteobacteria *Wolbachia* have evolved across the phyla nematoda and arthropoda. In *Wolbachia* phylogeny, supergroup F is the only clade known so far with members from both arthropod and filarial nematode hosts and therefore can provide unique insights into their evolution and biology. In this study, 4 new supergroup F *Wolbachia* genomes have been assembled using a metagenomic assembly and binning approach, *w*Moz and *w*Mpe from the human filarial parasites *Mansonella ozzardi* and *Mansonella perstans*, and *w*Ocae and *w*MoviF from the blue mason bee *Osmia caerulescens* and the sheep ked *Melophagus ovinus* respectively. A comprehensive phylogenomic analysis revealed two independent origins of filarial *Wolbachia* in supergroup F, most likely from ancestral arthropod hosts. The analysis also reveals that the switch from arthropod to filarial host is accompanied by a convergent pseudogenization and loss of the bacterioferritin gene, a phenomenon found to be shared by all filarial *Wolbachia*, even those outside supergroup F. These observations indicate that differences in heme metabolism might be a key feature distinguishing filarial and arthropod *Wolbachia*. The new genomes provide a valuable resource for further studies on symbiosis, evolution, and the discovery of new antibiotics to treat mansonellosis.

**Significance statement:** *Wolbachia* are bacterial endosymbionts of medically important parasitic filarial nematodes and arthropods. The evolutionary history and biological roles of *Wolbachia* in these different hosts are not well understood. The supergroup F in *Wolbachia* phylogeny harbors members from both filarial and arthropod hosts, providing an unparalleled opportunity for genomic comparisons to uncover distinguishing characteristics. This study provides new genomes from filarial and arthropod *Wolbachia* from this unique supergroup. Their phylogenomic analysis reveals multiple, independent transfers of *Wolbachia* from arthropod to filariae. Remarkably, these transfers were associated with a convergent loss of the bacterioferritin gene, a key regulator of heme metabolism. Heme supplementation is considered a critical component of *Wolbachia* - filaria symbiosis. We demonstrate bacterioferritin loss is a novel feature exclusive to filarial *Wolbachia*.

## Introduction

*Wolbachia* are gram-negative α-proteobacteria of the order *Rickettsiales*, present as intracellular symbionts in many species of parasitic filarial nematodes and arthropods (Werren et al. 2008). While the *Wolbachia* associations in arthropods range from reproductive parasites to nutritional mutualists (Werren et al. 2008; Zug & Hammerstein 2015), in filarial nematodes, *Wolbachia* is an obligate mutualist (Taylor et al. 2005) and essential for worm development, fecundity and survival (Taylor et al. 2005; Pfarr & Hoerauf 2006). *w*Mpe and *w*Moz are endosymbionts of *Mansonella perstans* and *Mansonella ozzardi*, respectively, the human filarial parasites responsible for mansonellosis. Mansonellosis is the most prevalent of all human filariases, yet the least studied and most neglected (Downes & Jacobsen 2010; Simonsen et al. 2011; Lima et al. 2016; Mediannikov & Ranque 2018; Ta-Tang et al. 2018; Sandri et al. 2021; Bélard & Gehringer 2021). *M. perstans* is considered to be the most common filaria in West and Central Africa (Simonsen et al. 2011; Ta-Tang et al. 2018), and is also found in the Amazon rainforest and the Caribbean coast of South America (Ta-Tang et al. 2018; Tavares da Silva et al. 2017). *M. ozzardi* infections have been reported from many countries in South and Central America, including some Caribbean Islands (Lima et al. 2016; Ta-Tang et al. 2018; Raccurt 2018; Calvopina et al. 2019). Co-infections of *M. perstans* and *M. ozzardi* are also known to exist in South America (Kozek et al. 1983; Crainey et al. 2020), complicating treatment as the two species respond differently to the commonly used anti-filarial drugs (Ta-Tang et al. 2018). Antibiotics targeting *Wolbachia* have been successful in treating human filarial diseases such as onchocerciasis and lymphatic filariasis (Pfarr & Hoerauf 2006; Langworthy et al. 2000; Bazzocchi et al. 2008; Clare et al. 2019; Taylor et al. 2019; Hübner et al. 2019; Hong et al. 2019, 10). Doxycycline has been shown to clear *M. perstans* microfilariae (Ta-Tang et al. 2018; Keiser et al. 2008; Coulibaly et al. 2009) and may have some efficacy against *M. ozzardi*, as well as dual infections with *M. ozzardi* and *M. perstans* (Crainey et al. 2020), however, no such clinical trials have been reported.

The *Wolbachia* present in the filarial nematodes of the genus *Mansonella* have a unique phylogenetic position compared to *Wolbachia* present in other human filarial parasites. The latest phylogenetic classification, based on Multi-Locus Sequence Typing (MLST), assigns *Wolbachia* to various monophyletic supergroups (Lefoulon et al. 2016; Lefoulon, Clark, Guerrero, et al. 2020; Lefoulon, Clark, Borveto, et al. 2020). Supergroups A, B, E, H and S are comprised entirely of arthropod *Wolbachia*, while supergroups C, D and J consist of filarial *Wolbachia* exclusively. Interestingly, the *Wolbachia* from *Mansonella* are classified under supergroup F, the only supergroup currently known to have *Wolbachia* members from filarial nematode as well as arthropod hosts. There is considerable interest in studying these *Wolbachia* to better understand the evolution and biology of arthropod and filarial *Wolbachia (*Lefoulon et al. 2016; Lefoulon, Clark, Guerrero, et al. 2020; Nikoh et al. 2014; Hosokawa et al. 2010).

Genomic information provides a fundamental resource for these investigations, yet only two genomes from supergroup F are currently available in public databases, namely the genomes of *Wolbachia w*Cle from the bedbug *Cimex lectularius* (Nikoh et al. 2014), and *w*Mhie, from *Madathamugadia hiepei*, a filarial parasite of geckos (Lefoulon, Clark, Guerrero, et al. 2020). In the current study, the genomes of supergroup F *Wolbachia*, *w*Moz and *w*Mpe from human filarial parasites *M. ozzardi* and *M. perstans* respectively, are assembled and analyzed. The genomes of arthropod-associated supergroup F *Wolbachia*, *w*Ocae and *w*MoviF, from the mason bee, *Osmia caerulescens,* and sheep ked, *Melophagus ovinus* are also reported. It is currently not known whether the *Wolbachia* of *M. ovinus* or *O. caerulescens* are facultative or obligate symbionts, or if they induce cytoplasmic incompatibility in these hosts. Genome-wide analyses of synteny, and annotations and comparative analysis of phage-derived gene content, cytoplasmic incompatibility genes and metabolic pathways of these genomes are described. In addition, comprehensive phylogenomic analyses of these 4 new genomes and all the available *Wolbachia* genomes were performed to generate comprehensive and robust phylogenetic trees, and confirmed the position of the new genomes within supergroup F. These analyses revealed at least two independent origins of filarial-*Wolbachia* association within supergroup F, most likely from ancestral arthropod-*Wolbachia* systems, and an accompanying loss of the bacterioferritin gene. The loss of bacterioferritin was discovered to be a feature common to all filarial *Wolbachia*.

## Results

### Genomes of *w*Moz and *w*Mpe, supergroup F *Wolbachia* from human filarial parasites *M. ozzardi* and *M. perstans*

A metagenomic assembly and binning pipeline suitable for complex clinical samples was utilized to analyze sequencing data from four *Mansonella* isolates. The binning of assembled metagenome scaffolds using BlobTools identified distinct clusters of scaffolds (“blobs”) corresponding to the *Wolbachia* genome, distinguishable from blobs corresponding to the genomes of the worm hosts and associated microbiota.

For the *M. ozzardi* isolate Moz1 from Brazil, only fragments of *Wolbachia* genome at low coverage were obtained and were not analyzed further (Supplementary Figure S1a). The BlobTools analysis of *M. ozzardi* isolate Moz2 from Venezuela (Supplementary Figure S1b) yielded a draft *w*Moz assembly of 1,073,310 bp in size comprised of 93 scaffolds. The N50 size of this assembly is 17.225 kb and the largest scaffold is 37.481 kb in length (Table 1). Of the two *M. perstans* isolates Mpe1 and Mpe2 from Cameroon, only Mpe1 yielded a *Wolbachia* assembly (Supplementary Figure S2). The *w*Mpe assembly is comprised of 1,058,123 bp in 170 scaffolds, with a N50 size of 10.041 kb and the largest scaffold 28.485 kb in length (Table 1).

**Table 1.** Characteristics of genomes of *wMoz* and *wMpe*, and other *Wolbachia* from the supergroup F

|  | ***w*Moz** | ***w*Mpe** | ***w*MoviF** | ***w*Ocae** | ***w*Mhie** | ***w*Cle** |
| --- | --- | --- | --- | --- | --- | --- |
| **Assembled genome size** | 1,073,310 bp | 1,058,123 bp | 1,008,858 bp | 1,201,397 bp | 1,025,329 bp | 1,250,060 bp |
| **Number of scaffolds** | 93 | 170 | 196 | 186 | 208 | 1 |
| **Largest scaffold** | 37.481 kb | 28.485 kb | 20.403 kb | 21.794 kb | 21.863 kb | - |
| **Scaffold N50 size** | 17.225 kb | 10.041 kb | 7.443 kb | 8.755 kb | 6.845 kb | - |
| **%GC** | 35.64 | 35.37 | 36.20 | 36.06 | 36.09 | 36.25 |
| **Predicted genes** | 1,079 | 1,058 | 1,009 | 1,173 | 1,081 | 1,246 |
| **Predicted proteins** | 888 | 864 | 928 | 1,055 | 907 | 981 |
| **Predicted pseudogenes** | 145 | 153 | 37 | 77 | 137 | 224 |
| **tRNA genes** | 36 | 34 | 35 | 34 | 30 | 34 |
| **BUSCO score (%)** | 78.1 | 75.3 | 81.3 | 80.4 | 79.0 | 81.7 |
| **Input reads** | This study | This study | ERR969522<br>Nováková et al. 2015 | SRR122170<br>Gerth et al. 2014 | Lefoulon et al. 2020 | Nikoh et al. 2014 |
| **Assembly accession and reference** | GCF_020278<br>625.1<br>This study | GCF_020278<br>605.1<br>This study | GCA_02366<br>1065.1<br>This study | GCA_02366<br>1085.1<br>This study | GCF_013366<br>855.1<br>Lefoulon et al. 2020 | GCF_000829<br>315.1<br>Nikoh et al. 2014 |

Whole genome alignment of *w*Moz and *w*Mpe demonstrated high sequence similarity and co-linearity of these independently assembled and closely related *Wolbachia* (Figure 1). A comparison was also made to the most closely related and the only complete genome available in the Clade F supergroup, namely *w*Cle, where high sequence synteny and co-linearity were also observed (Supplementary Figure S3).

**Fig. 1.**
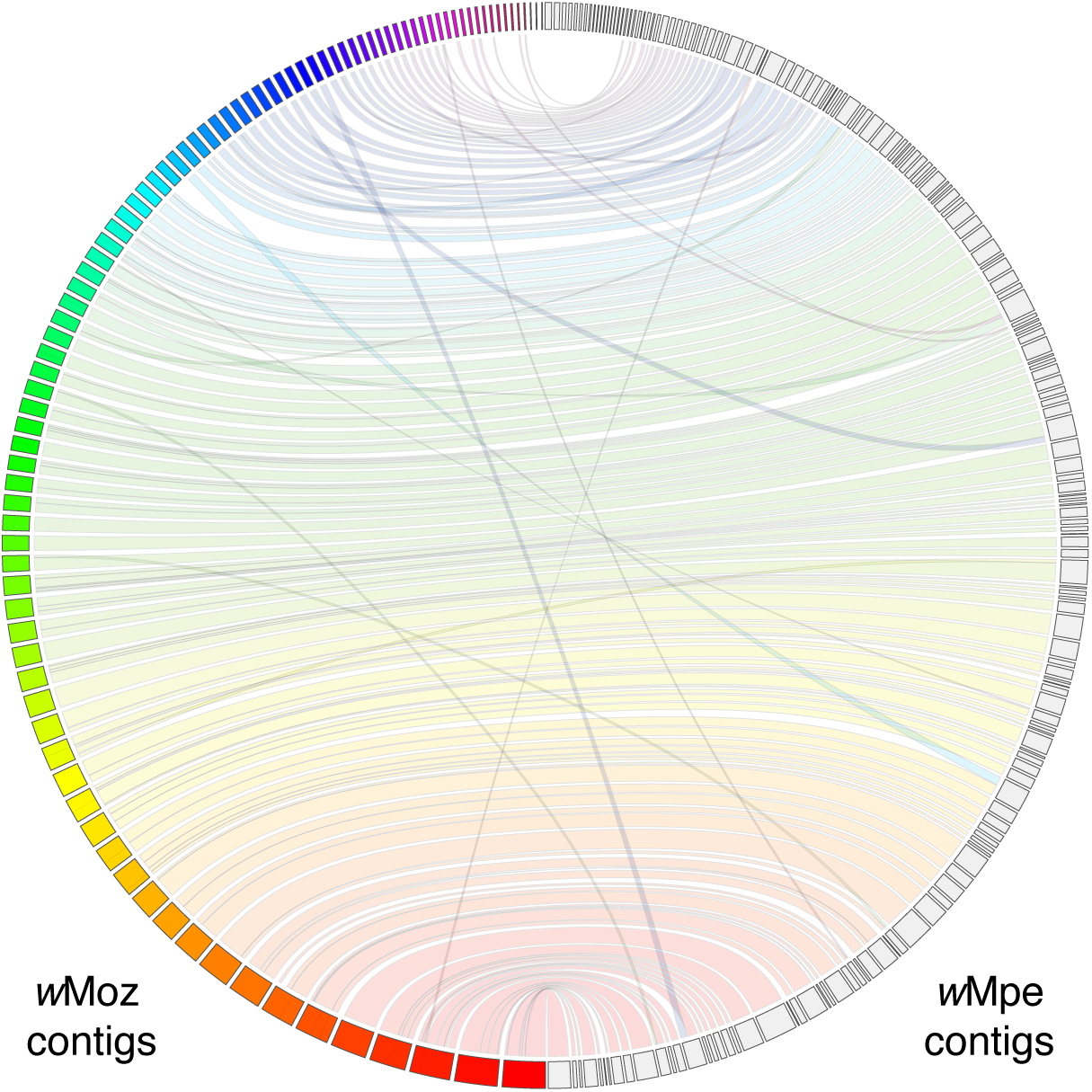
High sequence similarity and synteny between the *de novo* assembled *w*Moz and *w*Mpe genomes. The *Wolbachia* genomes assembled from two biologically and geographically distinct isolates display a high level of synteny and inter-genome consistency. The genomes were aligned using minimap2 and visualized using the JupiterPlot software. The *w*Moz scaffolds are displayed as colored boxes in the left semi-circle, while the *w*Mpe scaffolds are shown as gray boxes in the right semi-circle. The syntenic regions are marked by bands connecting scaffolds from one assembly to the corresponding region in the other assembly. Regions of rearrangement are shown as bands crisscrossing the horizontal bands of the syntenic regions.

Gene prediction using the NCBI PGAP pipeline identified 888 protein coding genes in *w*Moz and 864 protein coding genes in *w*Mpe. A BUSCO analysis (v5.1.3) was performed on these proteins to check for presence of the 219 genes conserved across most proteobacteria (“proteobacteria_odb10” database in BUSCO). The BUSCO scores of *w*Moz and *w*Mpe were 77.6% and 74.4% respectively (Table 1). These scores are typical for *Wolbachia,* even with complete genomes. For example, the BUSCO scores for *Wolbachia w*Ov from the human filarial parasite *Onchocerca volvulus*, *w*Oo from the bovine parasite *Onchocerca ochengi*, and *w*Bm from the human filarial parasite *Brugia malayi* are 76.7%, 75.8%, and 79.5% respectively (Supplementary Figure S4). Within supergroup F, the BUSCO score for filarial *Wolbachia w*Mhie is 79%, and it is 81.7% for the arthropod *Wolbachia w*Cle.

### Different isolates of *Mansonella* harbor varying levels of *Wolbachia*

The metagenomic assembly and binning approach enabled the determination of relative amounts of *Wolbachia* in different isolates by comparing read coverages of the *Mansonella-* and *Wolbachia-*specific scaffolds in the metagenomic assemblies (Table 2). In the *M. ozzardi* sample Moz2, the median coverage of scaffolds classified as *Mansonella* was 360X, while that of the *Wolbachia* scaffolds was 13X. Assuming 1000 nematode cells per microfilaria (Basyoni & Rizk 2016), each parasite is estimated to harbor 37 *Wolbachia* cells. For the other *M. ozzardi* isolate Moz1, the median coverage for nematode scaffolds was 57X and the corresponding *Wolbachia* coverage was only 3X. Therefore, titer calculations were not performed for the Moz1 isolate as they would not be robust at such low coverage and incomplete assembly. For the *M. perstans* sample (Mpe1), the median coverage of nematode scaffolds and *Wolbachia* scaffolds were 1,032X and 30X respectively, yielding an estimated titer of 30 *Wolbachia* per microfilaria. In the other *M. perstans* isolate Mpe2, even though the median coverage for nematode scaffolds is relatively high at 633X, only a very small portion of the *Wolbachia* genome, 15kb in 12 scaffolds (less than 2% of 1.073 Mb *w*Mpe1 genome) with a median coverage of 7X could be obtained during the metagenomic assembly, indicating a very low titer of *Wolbachia* in this isolate. The low titer was also confirmed using a complementary analysis, namely mapping the Mpe2 reads against the *w*Mpe1 assembly. In this analysis, the median depth of coverage obtained was only 9X, while the breadth of coverage (number of bases covered at 1X depth of coverage) was observed to be 83 kb only, just 8% of the *w*Mpe1 genome. Since both isolates were collected in Cameroon, these observations point to variation in *Wolbachia* titers within the same species in a geographical area and could explain the conflicting reports of PCR-based detection of *Wolbachia* in *M. perstans* (Sandri et al. 2021; Keiser et al. 2008; Coulibaly et al. 2009; Debrah et al. 2019; Grobusch et al. 2003; Gehringer et al. 2014).

**Table 2.**
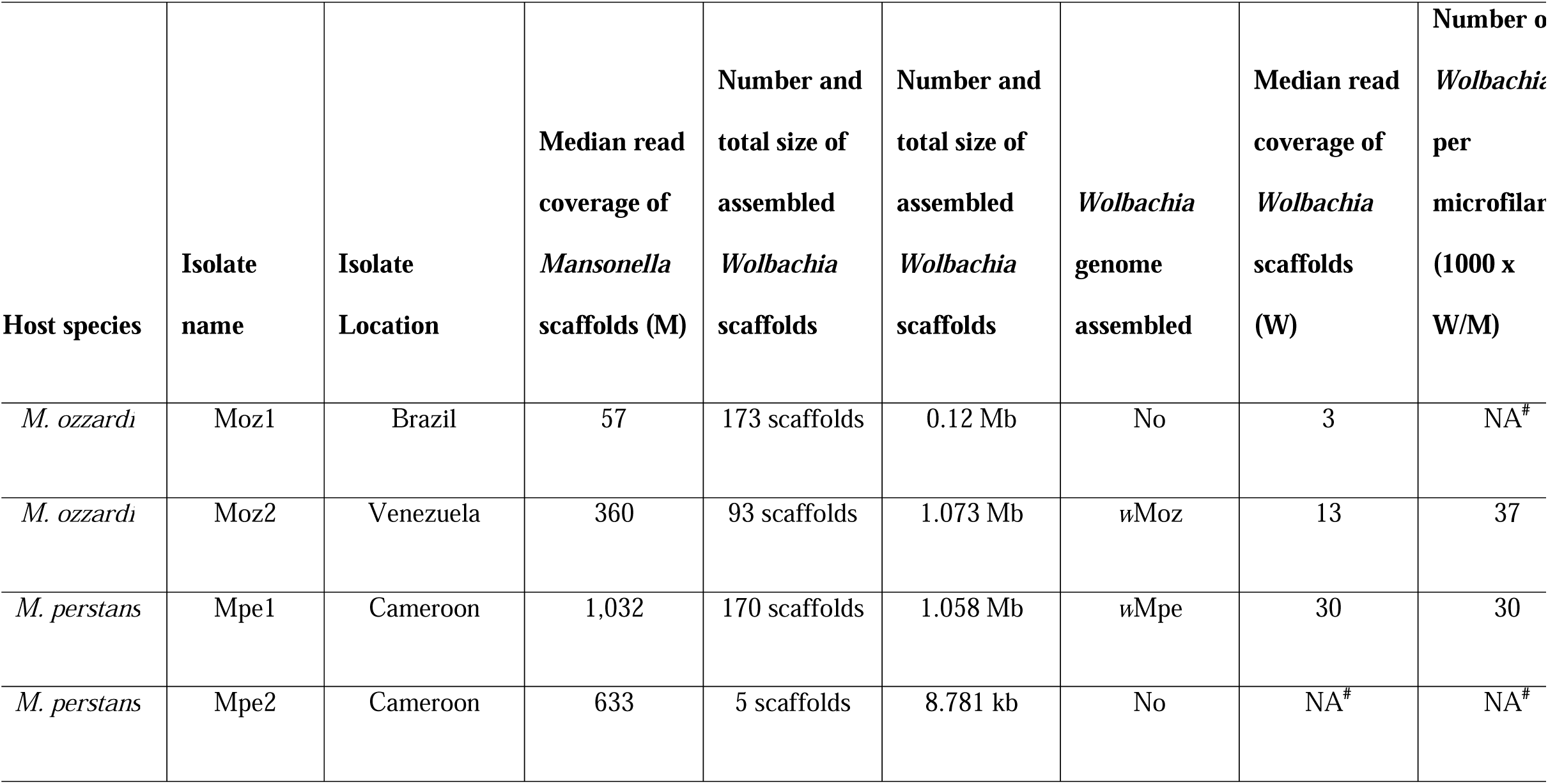
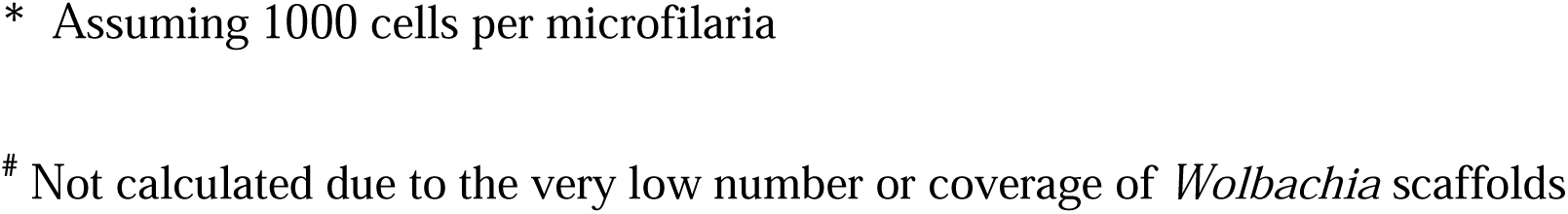
Estimation of *Wolbachia* levels per microfilaria

| **Host species** | **Isolate name** | **Isolate Location** | **Median read coverage of *Mansonella* scaffolds (M)** | **Number and total size of assembled *Wolbachia* scaffolds** | **Number and total size of assembled *Wolbachia* scaffolds** | ***Wolbachia* genome assembled** | **Median read coverage of *Wolbachia* scaffolds (W)** | **Number of *Wolbachia* per microfilaria (1000 x W/M)** |
| --- | --- | --- | --- | --- | --- | --- | --- | --- |
| *M. ozzardi* | Moz1 | Brazil | 57 | 173 scaffolds | 0.12 Mb | No | 3 | NA<sup>#</sup> |
| *M. ozzardi* | Moz2 | Venezuela | 360 | 93 scaffolds | 1.073 Mb | wMoz | 13 | 37 |
| *M. perstans* | Mpe1 | Cameroon | 1,032 | 170 scaffolds | 1.058 Mb | wMpe | 30 | 30 |
| *M. perstans* | Mpe2 | Cameroon | 633 | 5 scaffolds | 8.781 kb | No | NA<sup>#</sup> | NA<sup>#</sup> |

\* Assuming 1000 cells per microfilaria

### Genomes of *w*Ocae and *w*MoviF, supergroup F *Wolbachia* from the arthropods *O. caerulescens* and *M. ovinus*

For the *Wolbachia w*Ocae from the mason bee *O. caerulescens*, metagenomic assembly and binning was performed on Illumina sequencing reads available from a previous study (Gerth et al. 2014). A BlobTools analysis of the assembled scaffolds identified a distinct cluster of 186 *Wolbachia* scaffolds (Supplementary Figure S5a) with a total size of 1.2 Mb. A whole-genome alignment of these *w*Ocae contigs against the *w*Cle demonstrated high quality of this assembly (Supplementary Figure 5b). The NCBI-PGAP pipeline identified 899 protein coding genes, which had a BUSCO score of 79% (Table 1). During the preparation of this manuscript, a metagenomic assembly WOLB0012 of *Wolbachia* from the same raw data was published (Scholz et al. 2020), but only the un-annotated nucleotide sequence is available in NCBI database. A BlobPlot analysis of this assembly indicated the presence of arthropod contigs (Supplementary Figure S6). Therefore, instead of the WOLB0012 assembly, the carefully curated and annotated version of the *w*Ocae assembly described in this manuscript was used for all subsequent analyses.

The *Wolbachia* genome from the sheep ked *Melophagus ovinus* was obtained from analysis of publicly available reads. A first round of metagenomic assembly and BlobTools analysis revealed a complex mixture of multiple *Wolbachia* as well as other proteobacterial endosymbionts (Supplementary Figure S7). The total size of the 693 *Wolbachia* scaffolds identified in this analysis was 1.6 Mb, substantially larger than any *Wolbachia* genomes known so far. In addition, these scaffolds were spread over an unusually wide coverage range, over two orders of magnitude. A manual inspection of the blastn hits of this assembly revealed that some scaffolds had better hits against the genome of supergroup F *Wolbachia w*Cle, while others were more similar to the supergroup A *Wolbachia w*Mel or supergroup B *Wolbachia w*AlbB. Together, these observations point to multiple *Wolbachia* genomes in this sample. This complexity necessitated an iterative mapping and assembly approach utilizing only the reads that mapped to *Wolbachia* genomes, which produced 572 *Wolbachia* scaffolds with a total size 1.48 Mb (Supplementary Figure S8a). To identify the correct supergroup for each scaffold, its sequence similarity and query coverage in comparison to the *w*Cle, *w*Mel or *w*AlbB genomes were analyzed. The median percentage identity as well as query-coverage scores was observed to be the highest against the *w*Cle genome, followed by the *w*Mel genomes, and was much lower when compared to the *w*AlbB genome (Supplementary Figure S8b, 8c). Therefore, to isolate the supergroup F scaffolds, alignments of each scaffold to *w*Cle genome was compared to its alignments against the *w*Mel genome. Among the 572 scaffolds, 40 scaffolds had a nucmer hit only to the *w*Cle genome and were therefore included in the supergroup F bin, 214 scaffolds had a hit only to the *w*Mel genome (Supplementary Figure S8d) and were included in the supergroup A bin, while 84 scaffolds that did not have a hit in either *w*Cle of *w*Mel genome were found to be more similar to other *Wolbachia* from supergroups A or B. For the 234 scaffolds which had a nucmer hit to both *w*Cle and *w*Mel genomes, an affiliation score, calculated as the difference between its percentage identity to the *w*Cle genome and its percentage identity to the *w*Mel genome, was analyzed. This score showed a clear bimodal distribution, with a range from -15 to 13 (Supplementary Figure S8e) confirming the presence of two *Wolbachia* from different supergroups in the host *M. ovinus*. The scaffolds which had an affiliation score greater than zero were assigned to the supergroup F. These 156 scaffolds also showed a sequencing read-coverage distribution similar to the 40 scaffolds that had exclusive hits to the *w*Cle genome (Supplementary Figure S8f), confirming their classification into supergroup F. Finally, the classification of contigs as supergroup F was validated through an additional blobplot analysis using Illumina reads from a new metagenomic sequencing of a *M. ovinus* sample from China (Zhang et al. 2013). A combination of sequencing coverage from this new analysis and the affiliation scores described above achieved a clear classification (Supplementary Figure S8g). A whole-genome alignment of the contigs classified as *w*MoviF against the *w*Cle demonstrated high quality of this assembly (Supplementary Figure 8h). The final supergroup F bin comprised of 196 scaffolds with a total size of 1.01 Mb was designated as the genome for *Wolbachia w*MoviF, where the suffix F denotes the supergroup. NCBI-PGAP annotations identified 928 protein coding genes, which had a BUSCO score of 81.3% (Table 1) and no duplicated core genes (Supplementary Figure S4). The metagenomic bin for the putative supergroup A *Wolbachia* was highly fragmented (301 scaffolds, median length = 823 bp, total size = 0.387 Mb), and was not analyzed further. Recently, a metagenomic assembly of *Wolbachia* (GenBank accession WOLB1015), based on the same raw reads became available (Scholz et al. 2020). However, multiple analyses demonstrate that this assembly is a mixture of two different *Wolbachia*. First, the protein coding genes in WOLB1015 were predicted using PROKKA, and their BUSCO score was calculated. This analysis revealed duplications of 17% of single copy orthologs conserved across *Wolbachia*, an unusually high rate in contrast to 0% duplications that are observed in almost all other *Wolbachia* (Supplementary Figure S4). Second, a BlobPlot analysis of WOLB1015 showed the *Wolbachia* contigs spread over an unusually wide coverage range from 1X to 1000X, consistent with the presence of multiple *Wolbachia* in this assembly (Supplementary Figure S9). The WOLB1015 assembly had also failed to pass the QC criteria in the original study (Scholz et al. 2020). Therefore, instead of the mixed WOLB1015 assembly, the manually curated assembly of supergroup F *Wolbachia w*MoviF generated in this manuscript was used for all downstream analyses.

### Genome sequence comparisons between various *Wolbachia*

Analysis of whole genome alignments of the 4 newly assembled genomes to the complete *w*Cle genome showed that most of the regions absent from *w*Moz, *w*Mpe, *w*Ocae or *w*Mhie assemblies corresponded to regions containing IS element transposons in the *w*Cle genome (Figure 2). This indicates that most of the protein coding regions have been recovered in these draft genome assemblies, and the missing regions largely correspond to the IS elements, which due to their repetitive nature present a technical challenge to assemble using only short-read data (Sinha et al. 2019; Lefoulon et al. 2019).

**Fig. 2.**
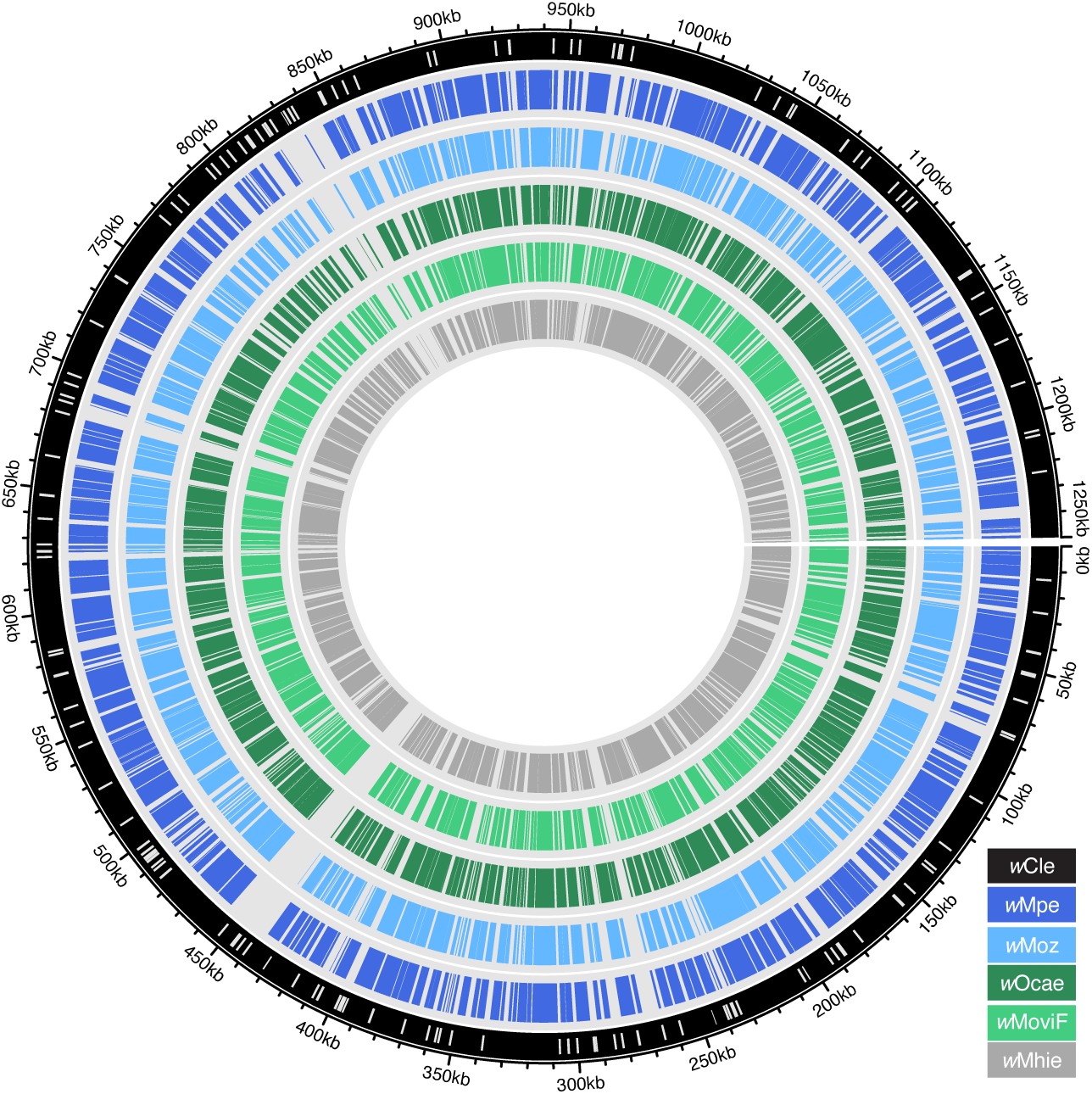
Gaps in the assembled *Wolbachia* genomes correspond mostly to IS elements. Whole genome alignments of *w*Mpe (blue ring), *w*Moz (light-blue ring), *w*Ocae (green ring), *w*MoviF (light green ring) and *w*Mhie (innermost gray ring) to the *w*Cle chromosome (outermost black circle) are visualized as a circos plot. The white bars mark the locations of the IS elements in the *w*Cle genome. The gaps in the alignments of various genomes to the *w*Cle genome overlap extensively with IS elements in the *w*Cle genome.

Comparisons of sequence similarity between all supergroup F *Wolbachia* genomes, and representatives from other supergroups including *Wolbachia* from filarial nematodes (Family Onchocercidae), a plant parasitic nematode *Pratylenchus penetrans* (Family Pratylenchidae), and arthropod hosts (Supplementary Table 1) were performed using genome-wide average nucleotide identity (gANI, Supplementary Figure S10a) and digital DNA–DNA hybridization (dDDH) scores (Supplementary Figure S10b-d). The dDDH scores provide more sensitivity when comparing closely related species (Meier-Kolthoff et al. 2013). The *w*Mpe:*w*Moz pair was found to have a dDDH score of 71.7% (Supplementary Figure S10b). For comparison, the dDDH scores for other closely related sympatric *Wolbachia* are: 96.1 % for *w*Oo:*w*Ov pair from *Onchocerca spp*., and 93.3 % for *w*Bm:*w*Bpa pair from *Brugia spp*. (Supplementary Figure S10c,d). These scores suggest *w*Mpe and *w*Moz have diverged substantially from each other as compared to the *w*Oo:*w*Ov or *w*Bm:*w*Bpa pairs, most likely due to them having split from their common ancestor before the *w*Bm:*w*Bpa or *w*Oo:*w*Ov split.

### Orthology analysis, gene function and pathway annotations

Protein-coding gene sequences from NCBI-PGAP annotations were obtained for all 123 *Wolbachia* genomes available in GenBank (Supplementary Table 1). For orthology analysis, protein-coding genes from the 4 new genomes, in addition to 77 genomes selected to represent all supergroups, were used (see column H, Supplementary Table 1). The OrthoFinder analysis of a total 82,406 proteins encoded by these 81 genomes identified 1,751 orthogroups, 99 of which represent Single Copy Orthologs (SCOs) conserved across all the analyzed *Wolbachia* (Supplementary Table 2). For the 4 new supergroup F *Wolbachia*, at least 96.9% of all predicted proteins could be assigned to an orthogroup (Supplementary Table 2).

Functional annotations of the proteins encoded by the *w*Moz, *w*Mpe, *w*MoviF and *w*Ocae genomes were obtained using the eggNOG database (Supplementary Table 3). Additionally, biosynthetic pathway annotations for the *w*Cle genome were obtained from the KEGG database, and corresponding orthologs were identified in the *w*Moz, *w*Mpe, *w*MoviF, *w*Ocae, and *w*Mhie proteomes. Genes encoding members of various metabolic pathways characteristic of *Wolbachia* proteomes, namely the heme pathway, purine and pyrimidine biosynthesis pathways, riboflavin metabolism, and Type IV secretion systems, were found to be mostly present in *w*Moz and *w*Mpe (Supplementary Table 4). Ribosomal protein subunits, a common target for anti-*Wolbachia* drugs such as doxycycline, as well as candidate drug targets such as pyruvate phosphate dikinase, PPDK (Raverdy et al. 2008), were also present in both *w*Moz and *w*Mpe.

### Biotin biosynthesis genes are absent from all supergroup F *Wolbachia* except *w*Cle

The bedbug *Wolbachia w*Cle genome harbors an operon encoding the enzymes involved in biosynthesis of biotin, which has been acquired via LGT from another *Rickettsia* (Nikoh et al. 2014). The biotin produced by *w*Cle supports a nutritional mutualism between the bedbug and its *Wolbachia* (Nikoh et al. 2014). To investigate whether this feature is common to other supergroup F *Wolbachia*, a search for the corresponding orthologs was performed. No orthologs of any of the genes involved in the biotin biosynthesis pathway (genes *bioA*, *bioD*, *bioC*, *bioH*, *bioF* and *bioB*) could be found in *w*Moz, *w*Mpe, *w*MoviF or *w*Ocae proteomes. Additionally, when the sequencing reads from these 4 supergroup F *Wolbachia* were mapped against the *w*Cle genome, no alignments could be found within the region harboring the biotin operon, even though alignments could be obtained in the neighboring regions flanking the biotin operon (Supplementary Figure S11). The absence of biotin pathway genes is unlikely to be due to the incomplete nature of the draft assemblies. Given the high BUSCO score (at least 79%) of the assemblies, if the biotin pathway was genuinely present in the genomes, at least fragments of a few of the 6 genes in the biotin pathway would have been detected. In contrast, the 7 genes for biosynthesis of riboflavin, another vitamin pathway important for the host-symbiont relationship (Li & Carlow 2012; Moriyama et al. 2015; Ju et al. 2020) were present in *w*Cle as well as all other supergroup F *Wolbachia* (Supplementary Table 4). Thus, biotin supplementation might be unique to the bedbug and its *Wolbachia* association but is not the basis of mutualism between other supergroup F *Wolbachia* and their corresponding filarial or arthropod hosts.

### Prophage remnants and mobile genetic elements in supergroup F *Wolbachia* genomes

Many *Wolbachia* are naturally infected with lambda-like temperate phages, termed WO phages, which can integrate into *Wolbachia* genomes as prophages (Bordenstein & Bordenstein 2022). The phage derived genes have been proposed to play important roles in generating genetic diversity, chromosomal evolution and transfer of genes required for *Wolbachia* symbiosis (Bordenstein & Bordenstein 2022). Phage-derived genes in all supergroup F *Wolbachia* were identified via a comprehensive search for phage-derived orthogroups (Figure 3, Supplementary Table 5) against a recently described database specific to *Wolbachia* phages (Bordenstein & Bordenstein 2022). In the *w*Moz and *w*Mpe genomes, 37 and 33 phage-derived orthogroups were found respectively, of which 32 were present in both *Wolbachia*. Among the gene members of these phage-derived orthogroups, “hypothetical protein” was the most common annotation (n = 22) followed by “ankyrin domain protein” (n = 12). The filarial *Wolbachia w*Mhie had 32 phage-associated orthogroups and shared 22 of these with *w*Moz and *w*Mpe. Neither *w*Moz, *w*Mpe, nor *w*Mhie were found to possess any core components of phage structural modules, like the filarial *Wolbachia* from other supergroups. Among arthropod *Wolbachia* of supergroup F, *w*Ocae had 29 phage structural genes, *w*MoviF had only 3 such genes, while *w*Cle had only one, suggesting different dynamics of phage evolution in these *Wolbachia*.

**Fig. 3.**
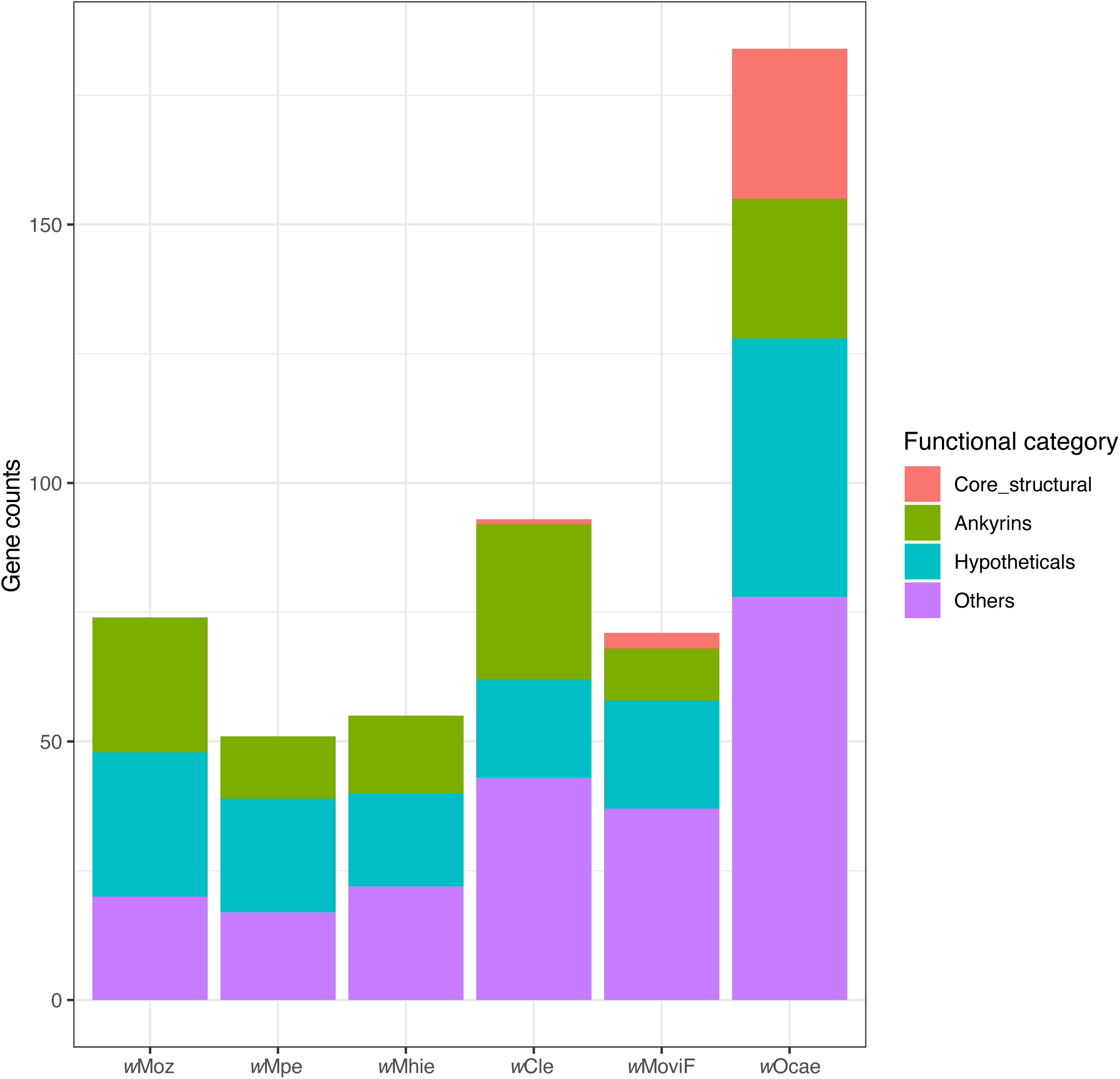
Numbers of phage-derived genes and their functional distributions in supergroup F *Wolbachia*. Phage derived genes in each *Wolbachia* were categorized according to their predicted protein function. Phage-derived genes which had annotations other than “Hypothetical protein” but could not be classified as either “Core structural” or “Ankyrins” were included in the category “Others”. Only *w*Ocae has intact phage modules, while *w*MoviF and *w*Cle have only one and two core structural phage genes respectively. Phage structural genes were absent from filarial *Wolbachia w*Moz, *w*Mpe and *w*Mhie.

Group II introns are common, phage-derived mobile elements (Bordenstein & Bordenstein 2022) that can be found in high copy numbers in many *Wolbachia* genomes: for example, 53 copies are found in the *w*AlbB genome (Sinha et al. 2019). Within the supergroup F genomes, the *w*Ocae genome was found to encode only two copies of Group II introns. No such genes could be detected in the *w*MoviF, *w*Moz or *w*Mpe genome assemblies. One pseudogene was detected in both *w*Moz and *w*Mpe. No intact gene or pseudogene for these elements could be found in the *w*Mhie or *w*Cle genomes, as reported earlier (Lefoulon, Clark, Guerrero, et al. 2020).

The *Wolbachia cifA*-*cifB* genes, which encode cytoplasmic incompatibility factors that play a key role in the reproductive biology of their arthropod hosts, are also derived from phages (Bordenstein & Bordenstein 2022; Lindsey et al. 2018). In supergroup F, no *cifA* or *cifB* genes could be detected in the genomes of arthropod *Wolbachia w*Cle or *w*MoviF. In *w*Ocae, multiple pseudogenes or fragments of *cifA* and *cifB* genes missing either the 5-prime or 3-prime ends were identified (Supplementary Table 6). Since all the gene fragments were observed to be located at the termini of the assembled scaffolds, they are most likely to be pseudogenes disrupted by IS element insertions that tend to fragment the assemblies (Sinha et al. 2019). Thus, *w*Ocae also seems to be devoid of any intact and functional *cifA*-*cifB* gene pair. No *cifA* or *cifB* orthologs could be detected in filarial *Wolbachia w*Moz, *w*Mpe, or *w*Mhie.

### Phylogenomic analysis encompassing all *Wolbachia* genomes

To determine the evolutionary relationships between the supergroup F *Wolbachia* and other *Wolbachia*, as well as within supergroup F, comprehensive phylogenomic analyses were performed. Using a total of 127 *Wolbachia* genomes, including all 123 annotated genomes available from GenBank (Supplementary Table 1) as input for OrthoFinder, 46 Single Copy Orthologs (SCOs) that are conserved across all available genomes were identified. The corresponding phylogenetic tree based on their multiple sequence alignment spanning 8,394 amino acids was produced (Supplementary Figure S12). For a further refinement of the tree, all incomplete or redundant genomes were excluded from analysis, except the genomes within supergroup F (see column H, Supplementary Table 1). OrthoFinder analysis on these selected 81 genomes identified 99 SCOs (Supplementary Table 2). Maximum likelihood trees with a partitioned model were built from the combined supermatrix of 62,160 nucleotides (Figure 4) and 20,470 amino acids (Supplementary Figure S13). The 4 new genomes were robustly placed within supergroup F in this analysis. In this tree, the *Wolbachia w*CfeJ from the cat flea *C. felis*, which has not been assigned any supergroup yet, was found to be closest to supergroup F.

**Fig. 4.**
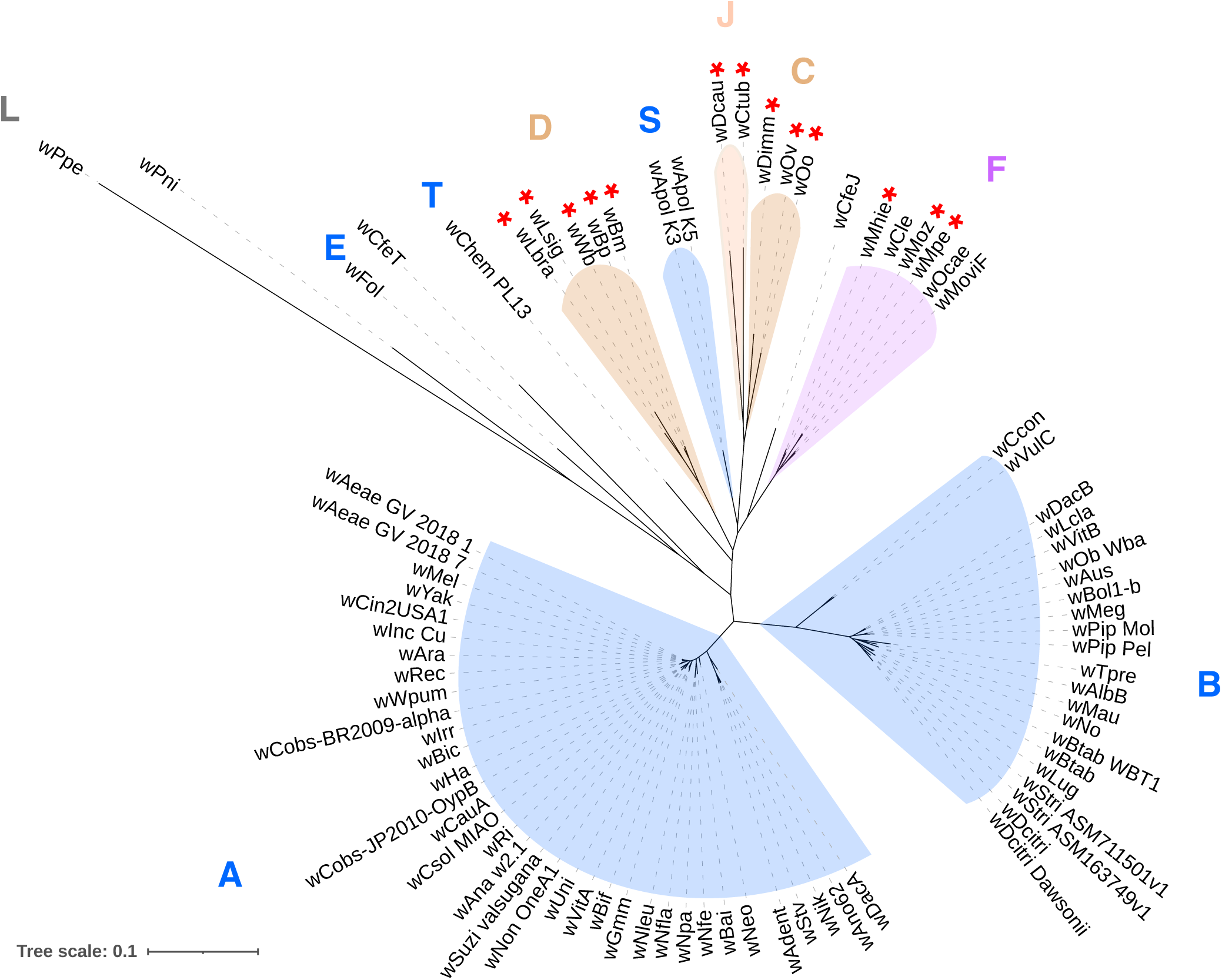
Global phylogenomic analysis of all *Wolbachia* genomes. Phylogenomic tree based on gene sequences of 91 single copy orthologs (SCOs) conserved across 81 *Wolbachia* representing all supergroups. Supergroups comprised of *Wolbachia* exclusively from arthropod hosts are marked in blue, while the supergroups comprised exclusively of filarial *Wolbachia* are marked in orange. The supergroup F, which has members from both arthropod and filarial hosts, is marked in purple background. The red asterisks denote the *Wolbachia* with a pseudogenized or absent bacterioferritin gene.

Remarkably, the filarial *Wolbachia w*Moz and *w*Mpe were placed in a clade different from the clade harboring *w*Mhie, the other filarial *Wolbachia* in this supergroup (Figure 5A). The filarial *Wolbachia w*Moz and *w*Mpe were most closely related to the arthropod *Wolbachia w*MoviF and *w*Ocae, while the filarial *Wolbachia w*Mhie was more closely related to the arthropod *Wolbachia w*Cle. Given the robust bootstrap support values for the branches (Figure 5A), the most likely recent common ancestor for the *w*Moz, *w*Mpe lineage would be an arthropod *Wolbachia* closer to *w*MoviF and *w*Ocae, while the *w*Mhie and *w*Cle clade would have received their *Wolbachia* from a different arthropod-*Wolbachia* ancestral donor. The topology of the *Wolbachia* phylogeny was also compared to the phylogeny of multiple *Wolbachia* hosts (Figure 5A), and no congruence was found between the two phylogenies (Figure 5A). Particularly, within supergroup F, the clear separation of the filarial *Wolbachia* into two separate, well-supported clades, and the discordant phylogenetic topology of their corresponding hosts reveals that the filaria-*Wolbachia* symbiosis has originated at least twice within this supergroup, from a different ancestral arthropod-*Wolbachia* donor in each case.

**Fig. 5.**
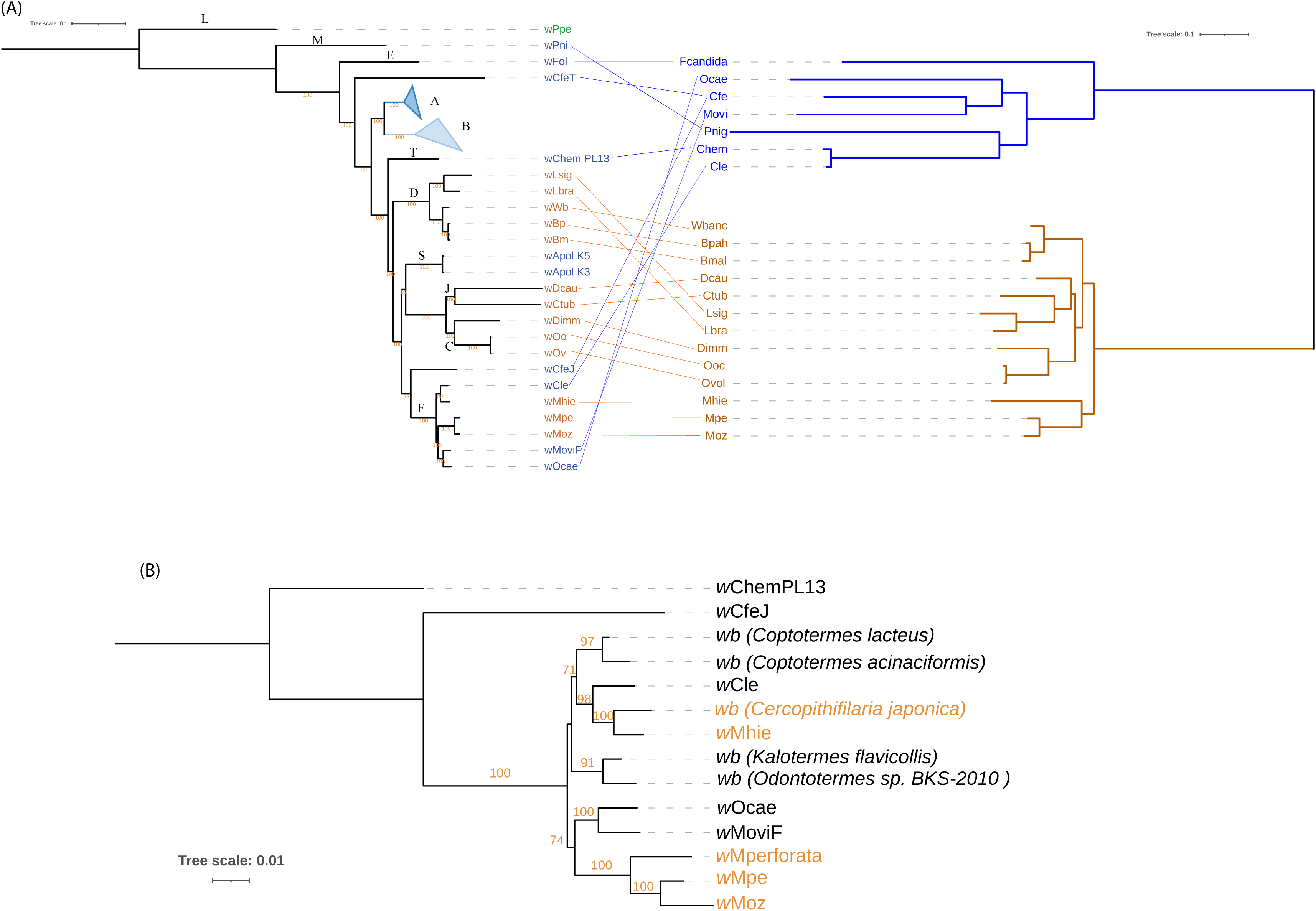
Two independent origins of *Wolbachia*-filaria symbiosis within supergroup F. (a) Mid-point rooted phylogenomic tree of *Wolbachia* based on gene sequences of 91 SCOs conserved across *Wolbachia* juxtaposed with a phylogenetic tree of various *Wolbachia* hosts based on 1,336 orthogroups. All filarial species, their *Wolbachia* and the lines depicting host-*Wolbachia* relationships are in orange The arthropod hosts, their *Wolbachia* and their connecting lines are labelled in blue. The bootstrap values are shown along the respective branches. Supergroups A and B comprise of 35 and 22 *Wolbachia* genomes respectively and are represented together as blue triangles. *Wolbachia* supergroups are denoted near the root node of each supergroup. (b) Phylogenetic tree of supergroup F *Wolbachia* based on a supermatrix of 8 genes (*ftsZ*, *fbpA*, *coxA*, *gatB*, *hcpA*, *groEL*, *dnaA*, 16S rRNA), total sequence length 10,374 nucleotides, also supports at least two independent origins of filarial *Wolbachia*. All *Wolbachia* from *Mansonella* hosts, including *M. perforata* are placed in a clade adjacent to the clade harboring arthropod *Wolbachia w*Ocae and *w*MoviF, while the *Wolbachia w*Mhie and *w*Cjaponica from filarial hosts *M. hiepei* and *Cercopithifilaria japonica* are in a separate clade that is more closely related to the arthropod *Wolbachia w*Cle. Filarial *Wolbachia* are marked with orange font. Only the standard bootstrap values higher than 70 are shown.

To expand the phylogenetic analysis of supergroup F *Wolbachia* beyond those with whole genome sequences, gene sequences for 8 genes, including all MLST loci, as well as *groEL*, *dnaA*, and 16S rRNA genes were collected from GenBank (Supplementary Table 7). The corresponding sequences from *w*CfeJ and *w*ChemPL13 were used as outgroups. This tree (Figure 5B) also robustly supports the separation of the *Mansonella Wolbachia* clade from the *w*Mhie clade. Interestingly, the only other known filarial *Wolbachia* in supergroup F, from the deer tick nematode *Cercopithifilaria japonica*, is placed in a clade with *w*Mhie, with the arthropod *Wolbachia w*Cle sharing a common ancestor with both these filarial *Wolbachia*.

### Convergent loss of bacterioferritin across filarial *Wolbachia*

Genome reduction and gene loss is a recurrent theme in the evolution of endosymbionts (Andersson & Kurland 1998), and the genomes of filarial *Wolbachia* are often found to be even more reduced as compared to their arthropod counterparts. A study of the genes lost during this evolutionary transition can therefore reveal genes that are either potentially dispensable or have been selected against during the evolution of filaria-*Wolbachia* symbiosis. The analysis of orthology relationships across 81 *Wolbachia* genomes (Supplementary Table 2) identified various orthogroups that are absent in filarial *Wolbachia* but are present in arthropod *Wolbachia*. This approach uncovered a striking loss of the bacterioferritin gene in filarial *Wolbachia* across all supergroups C, D, J and F (Figure 4). Within supergroup F, at least two independent losses of the bacterioferritin gene were observed, one in the *Mansonella Wolbachia* clade and the other in the *w*Mhie clade (Figure 5a), with a missing canonical start codon and premature stop codon resulting in a truncated protein about 75% of the length of the intact protein (115 aa in *w*Mpe, 125 aa in *w*Moz, 121 aa in *w*Mhie as compared to 160 aa in *w*Cle). The bacterioferritin proteins functions as a complex of 24 subunits forming a spherical cage to trap heme molecules (Rivera 2017) and a truncated protein is likely to disrupt this complex structure. Within supergroup F, all filarial *Wolbachia*, namely *w*Moz, *w*Mpe and *w*Mhie, contain a pseudogenized bacterioferritin, even though their most closely related arthropod *Wolbachia* have the corresponding gene intact. Since the *w*Moz:*w*Mpe pair is placed in a separate clade from *w*Mhie (Figure 5), our results suggest that the *bfr* gene has been pseudogenized independently in the two separate lineages.

Interestingly, pseudogene fragments of the *bfr* gene in various stages of degradation could be found across filarial *Wolbachia* (Supplementary Figure S15a). In supergroups C and J, although *bfr* pseudogenes were not found in the NCBI-PGAP annotations, blastx searches could identify leftover fragments of bacterioferritin in the corresponding genomic region (Supplementary Figure S15a). This genomic region was observed to be syntenic across all analyzed genomes (Supplementary Figure S15a), confirming that the lack of the *bfr* gene is not due to assembly artefacts.

Although the bacterioferritin locus in the RefSeq annotations of *Wolbachia w*Bpa from *Brugia pahangi* is annotated in NCBI as an intact CDS instead of a pseudogene, a closer look at its sequence and multiple sequence alignment with other bacterioferritin genes points to the loss of the canonical start codon ATG, similar to the corresponding sequence in *w*Bm, as well as other nucleotide insertions and deletions downstream (Supplementary Figure S15b), suggesting the pseudogenization of this gene in *w*Bpa as well. In *w*Bm, the *bfr* locus has a premature stop codon resulting in truncated protein with 138 amino acids instead of the complete 160 amino acids. The predicted peptide sequences corresponding to these pseudogenes showed multiple premature stop codons and frameshifts in both *w*Bm and *w*Bpa (Supplementary Figure S15c).

The dN/dS ratio, the ratio of non-synonymous to synonymous substitutions for the *bfr* locus in *w*Bpa was found to be higher than the corresponding locus in *w*Bm, (Supplementary Table 9). Prior to dN/dS calculations, analysis of transitions and transversions with respect to genetic divergence plots and calculation of substitution saturation index (Xia et al. 2003) was performed to ensure that the bacterioferritin locus has not undergone substitution saturation across the *Wolbachia* (Supplementary Figure S16). Further, the dN/dS values for *bfr* locus in *w*Bm, *w*Bp and were also compared against the distribution of genome-wide dN/dS ratios for all potential pseudogenes detected in these genomes using Pseudofinder (Syberg-Olsen et al. 2022). This analysis showed the dN/dS values for the *bfr* locus in *w*Bm and *w*Bpa are substantially higher than the median values of genome-wide dN/dS (Supplementary Figure S17). Together, these analyses confirm that bacterioferritin is undergoing pseudogenization in all filarial *Wolbachia*.

### Analysis of pseudogenes shared within the supergroup F *Wolbachia*

Intracellular bacterial symbionts, such as *Wolbachia*, are expected to accumulate pseudogenes due to genetic drift over evolutionary timescales (Andersson & Kurland 1998). However, the observation of two independent *bfr* pseudogenization events within supergroup F *Wolbachia* suggests that additional factors, such as convergent evolution through negative selection on certain genes could play a role in pseudogene formation. To explore the contributions of these factors, pseudogenes shared between various supergroup F *Wolbachia* were analyzed. This involved first determining the orthogroup corresponding to the ancestral gene of each pseudogene (Supplementary Table 8). These “pseudogene-orthogroups” were then compared to identify those unique or shared across different *Wolbachia* (Figure 6). For each *Wolbachia*, the majority of its pseudogene-orthogroups were found to be unique to its genome, whereas typically very few (n = 1 to 6) were shared between different *Wolbachia*, consistent with the genetic drift model. An exception to this trend was the number shared between the filarial *Wolbachia w*Moz, *w*Mpe and *w*Mhie (n = 15, Figure 6), which included the bacterioferritin pseudogene described above. Since *w*Moz and *w*Mpe are placed in a clade separate from *w*Mhie, these 15 pseudogene-orthogroups have most likely arisen from independent, convergent loss in filarial *Wolbachia* across different lineages within supergroup F. Since the related arthropod *Wolbachia* all have intact counterparts for each of these 15 genes, arthropod *Wolbachia* are the most likely ancestral sources for these filarial *Wolbachia* in supergroup F. The highest number of shared pseudogene-orthogroups was observed in the *w*Moz:*w*Mpe pair (n = 39). The corresponding pseudogenes might have arisen either through convergent evolution specific to *Wolbachia* within *Mansonella* or were formed by genetic drift in the last common ancestor of *w*Moz and *w*Mpe, or a combination of both factors.

**Fig. 6.**
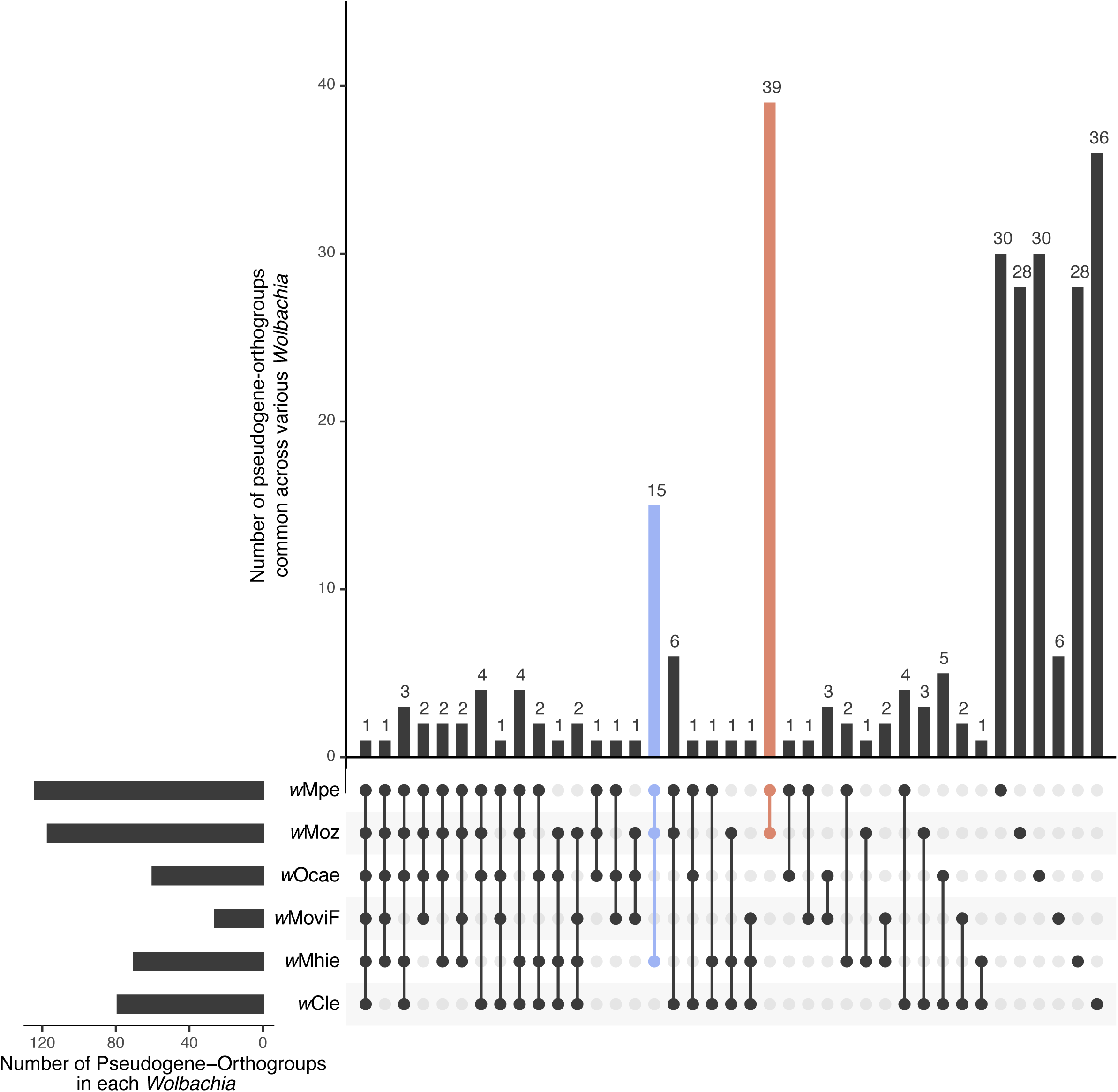
Pseudogenes shared between supergroup F *Wolbachia*. The orthogroups corresponding to the ancestral gene of each pseudogene were compared across different *Wolbachia*. Each *Wolbachia* harbors its unique set of pseudogenes, reflecting their origin as evolutionary accidents. The largest set of shared pseudogenes is observed between the closely related *w*Moz and *w*Mpe *Wolbachia* (n = 39, dark orange bar), suggesting either a shared common ancestor or convergent evolution within their respective *Mansonella* hosts. The next largest shared set of pseudogenes is found among the *Wolbachia w*Moz, *w*Mpe, and *w*Mhie (n = 15, blue bar), which are phylogenetically in different clades, indicating that the corresponding genes have undergone convergent loss of function across the three filarial *Wolbachia* in supergroup F. Transposons such as IS elements were excluded from this analysis due to their tendency to frequently convert into pseudogenes.

## Discussion

*Wolbachia* endosymbionts play critical roles in the biology of their arthropod and filarial hosts, and there is great interest in understanding their diverse functions through genomic approaches. The enigmatic supergroup F in *Wolbachia* phylogeny is the only supergroup comprising symbionts of arthropod as well as filarial nematode hosts. In this report, 4 new *Wolbachia* genomes from supergroup F are described, expanding the number of genomes from 2 to 6 in this under-represented supergroup. Two genomes are from *w*Moz and *w*Mpe, endosymbionts of Mansonella *spp.*, important filarial parasites of humans, while the other 2 genomes are from *w*Ocae and *w*MoviF, endosymbionts of the mason bee *O. caerulescens* and the sheep ked *M. ovinus* respectively.

Since *Wolbachia* are intracellular symbionts that cannot be cultured independently of their hosts, it is not possible to obtain their DNA independent of the host and the associated microbiome, presenting a challenge for accurate genome assembly. In the case of *w*Moz and *w*Mpe, complex human clinical samples were used as the source material, containing a mixture of DNA from human blood, the filarial parasite host, its intracellular symbiont *Wolbachia*, as well as other constituents of the human microbiome. Similarly, for *w*Ocae and *w*MoviF genomes, the publicly available sequencing data was derived from field-collected, wild isolates of their arthropod hosts, and comprised DNA from the arthropod host, *Wolbachia*, as well as the arthropod microbiome. Therefore, a metagenomic assembly and binning pipeline, customized to each of the datasets, was necessary and played a critical role in obtaining high quality *Wolbachia* genomes.

The BlobTools-based binning approach separates the assembled scaffolds based on their GC content and relative read coverage, which also provides a means for an accurate determination of *Wolbachia* titers via the measurement of relative copy numbers of genomes of *Wolbachia* versus its host. This principle has been used in a metagenomic study for *Wolbachia* titer calculations across multiple hosts (Scholz et al. 2020), and for highlighting sex-specific differences in *Wolbachia* levels in *D. immitis* (Kumar et al. 2013). Given the longstanding controversy around the detection of *Wolbachia* in *M. perstans* (Sandri et al. 2021; Keiser et al. 2008; Coulibaly et al. 2009; Debrah et al. 2019; Grobusch et al. 2003; Gehringer et al. 2014), determination of *Wolbachia* titers in *Mansonella* is particularly important, and has implications for antibiotic use as a treatment (Gehringer et al. 2014). In the current study, the titer in the isolate Mpe1 was found to be 30 *Wolbachia* cells per microfilaria. In the isolate Mpe2, although the presence of *Wolbachia* sequences could be detected, their depth and breadth of coverage was too low for estimating the titer reliably. Importantly, this low *Wolbachia* coverage in Mpe2 is not simply due to an overall low sequencing depth, as the corresponding genome for the host *M. perstans* could be successfully assembled at a high coverage that was comparable to the isolate Mpe1. As both Mpe1 and Mpe2 samples were collected in Cameroon, these observations demonstrate that even within the same geographical region, *Wolbachia* levels can widely vary between different isolates. For the *M. ozzardi* isolate Moz1, *Wolbachia* was detected but could not be assembled due to overall low sequencing depth obtained for this sample. In isolate Moz2, the titer was determined to be 37 *Wolbachia* cells per microfilaria, similar to Mpe1, and to levels reported from a laboratory strain of *Brugia malayi* (McGarry et al. 2004). The presence of *Wolbachia* in all *Mansonella* isolates described here, albeit at varying levels, and in another human parasite *Mansonella* spp. “DEUX” (Sandri et al. 2021), together with the successful clearance of *M. perstans* microfilariae with doxycycline (Coulibaly et al. 2009; Debrah et al. 2019), supports the use of antibiotics as a promising treatment for mansonellosis.

The *Wolbachia* from supergroup F present a unique opportunity to investigate the evolutionary history of closely related arthropod and filarial *Wolbachia* since all other supergroups are comprised exclusively of either arthropod or filaria associated *Wolbachia*. A global phylogenomic analysis of 127 *Wolbachia* genomes, including all supergroup F genomes, confirmed the placement of the 4 new genomes (*w*Moz, *w*Mpe, *w*MoviF, and *w*Ocae) in supergroup F, and identifies *w*CfeJ as the most closely related and robust outgroup to this supergroup. The phylogenomic analyses revealed two independent origins of filarial *Wolbachia* within supergroup F. The *w*Moz and *w*Mpe pair shared a common ancestor with the arthropod *Wolbachia w*Ocae and *w*MoviF, while *w*Mhie shared a different common ancestor with the bed bug *Wolbachia w*Cle. While these evolutionary scenarios supporting multiple horizontal transfers of *Wolbachia* into filaria have previously been proposed, they were based on phylogenetic studies utilizing sequences from as few as 2 to 7 gene loci (Casiraghi et al. 2005; Lefoulon et al. 2012, 2016). The genome-wide phylogeny in this study is based on a supermatrix of 91 genes, and incorporates 2 new filarial and 2 new arthropod *Wolbachia* genomes, thereby providing the most comprehensive and robust evidence in support of the independent origins. The widespread occurrence and transmission of *Wolbachia* infections across different animal phyla can occur through cladogenic inheritance, introgression or horizontal transfer. The relative contributions of three transmission modes to evolutionary trajectories have been analyzed in detail for *Wolbachia* spread in closely related host systems such as in the *Nasonia* species complex (Raychoudhury et al. 2009) and *Drosophila* species complex (Turelli et al. 2018; Cooper et al. 2019), and for reproductively isolated *Drosophila* species *D. simulans* and *D. ananassae*, the horizontal transfer of *Wolbachia* was found to explain the observed phylogeny (Turelli et al. 2018). Since, the arthropod and filarial hosts in supergroup F are also from widely diverged and reproductively isolated species and phyla, horizontal transfer of *Wolbachia* between the two phyla is the most likely explanation for their observed phylogeny. This conclusion is further supported by the incongruence between the phylogenies of *Wolbachia* and their hosts.

The phylogenomic analysis also reveals the complex, interleaved pattern of filaria-only supergroups C, D and J and arthropod-only supergroup S and the arthropod *Wolbachia w*CfeJ. In the current phylogeny, the supergroup T *Wolbachia w*ChemPL13 from bed bug is observed to be basal to all these supergroups, with robust bootstrap support. In principle, it is still possible that a currently unknown nematode *Wolbachia* was the ancestral state, but the current phylogenetic position of *w*ChemPL13 and nesting patterns of other subgroups favors the model where filarial *Wolbachia* are most likely acquired form an arthropod-associated ancestor.

Additional evidence for the direction of *Wolbachia* transfer being from arthropod to filarial hosts in supergroup F is provided by the striking observation of a convergent pseudogenization of the bacterioferritin gene in filarial *Wolbachia*. Within supergroup F, at least two independent losses of the bacterioferritin gene were observed, one in the *w*Moz-*w*Mpe clade and the other in the *w*Mhie clade, while their sister *Wolbachia* from arthropod hosts had an intact *bfr* gene, implying the intact *bfr* gene and therefore arthropod *Wolbachia* as the ancestral state.

The loss of bacterioferritin has important implications for the obligate relationship between *Wolbachia* and filarial hosts. Bacterioferritin is known to be a critical regulator of heme homeostasis in bacteria, serving as a store for heme that can be used during heme deficiency, or sequestering excess heme to prevent toxicity (Carrondo 2003). In arthropods, the presence of *Wolbachia* provides a fitness benefit by increasing fecundity during conditions of iron deficiency (Brownlie et al. 2009), while during iron excess, *Wolbachia* protect their insect hosts from iron induced toxicity by upregulating bacterioferritin (Kremer et al. 2009). Heme is an essential nutrient required by filarial worms, but they lack a heme biosynthetic pathway. Heme provisioning by *Wolbachia* has been proposed to play a key role in the obligate host-symbiont relationship (Foster et al. 2005; Gill et al. 2014). The lack of bacterioferritin in filarial *Wolbachia* suggests that *Wolbachia* cells can produce but no longer store heme intracellularly, making it available for uptake by the host filaria, and in doing so, affording a fitness advantage for the parasite. Under this model, the various ancestral filarial *Wolbachia* might have had an intact bacterioferritin gene to begin with, but in the lineages that lost this gene due to genetic drift, the freeing up of heme for the filarial host would have provided a strong enough fitness advantage to drive the *bfr*-lacking *Wolbachia* to fixation in filarial hosts and initiating an obligate relationship. Although genome reduction and gene loss, usually via genetic drift, is a recurrent theme in the evolution of endosymbionts (Andersson & Kurland 1998), the loss of bacterioferritin gene across multiple filarial *Wolbachia* is a unique example of convergent gene loss accompanying the adaptation to a new ecological niche. Further research into the role of bacterioferritin and heme metabolism can further elucidate their role in different host-*Wolbachia* systems.

In summary, the comprehensive analysis of newly assembled supergroup F *Wolbachia* genomes have provided the first genomic support for multiple origins of filaria and *Wolbachia* symbiosis from ancestral arthropod hosts and uncovered an intriguing loss of bacterioferritin from all filarial *Wolbachia*. Further studies of these genomes are expected to shed more light on the evolution and nature of symbiosis, as well as the identification of new treatments for mansonellosis.

## Materials and Methods

### Ethics statement

All research involving human subjects was approved by the appropriate committees and performed in accordance with all relevant guidelines and regulations. Informed consent was obtained from all participants or their legal guardians.

For *M. perstans*, ethical clearance was obtained from the National Institutional Review board, Yaoundé (REF: N° 2015/09/639/CE/CNERSH/SP) and administrative clearance from the Delegation of Public Health, South-West region of Cameroon (Re: R11/MINSANTE/SWR/ RDPH/PS/259/382). Approval for the study was granted by the “National Ethics Committee of Research for Human Health” in Cameroon. Special consideration was taken to minimize the health risks to which any participant in this study was exposed. The objectives of the study were explained to the consenting donor after which they signed an informed consent form. The participant’s documents were assigned a code to protect the privacy of the study subject. At the end of the study, the donor received a cure of mebendazole (100 mg twice daily for 30 days).

For *M. ozzardi*, study protocols were approved by the Institutional Review Board of the Institute of Biomedical Sciences, University of São Paulo, Brazil (1133/CEP, 2013). Written informed consent was obtained from all patients, or their parents or guardians if participants were minors aged <18 years. Diagnosed infections were treated with a single dose of 0.2 mg/kg of ivermectin after blood sampling (Basano et al. 2018; Lima et al. 2018, 39).

### Parasite materials and DNA extraction

*M. ozzardi* DNA was prepared from microfilariae collected from blood samples of individuals who participated in a previous study in Brazil (Lima et al. 2018). A sample containing a high microfilaria load was selected for genome sequencing and is denoted as Moz1 in the present study. A Venezuelan isolate of *M. ozzardi* microfilariae was generously donated by Izaskun Petralanda in 1989 and is denoted as Moz2 in this study. Genomic DNA was prepared from Moz2 microfilariae by Proteinase K digestion followed by phenol/chloroform extraction and ethanol precipitation as well as drop dialysis (https://www.neb.com/protocols/2013/09/16/drop-dialysis) then stored at –20°C. Two independent isolates of *M*. *perstans* microfilariae, denoted as Mpe1 and Mpe2 in this study, were collected on nylon filters from blood samples from Cameroon (Poole et al. 2019). Mpe1 DNA was extracted using a Genomic DNA Tissue MicroPrep kit (Zymo Research, USA) as described previously (Poole et al. 2019). DNA from Mpe2 was isolated as described above for Moz2. DNA quantity was determined using a Qubit dsDNA HS Assay kit in conjunction with a Qubit 2.0 Fluorometer as directed by the manufacturer (Life Technologies, USA).

### Illumina library construction and sequencing

Prior to library construction, the NEBNext Microbiome DNA enrichment kit (New England Biolabs Inc., USA) was used to enrich parasite DNA and reduce human DNA contamination (except for the Moz2 sample). The enrichment process is based on selective binding and removal of CpG-methylated DNA islands present in human genome but absent from nematode and *Wolbachia* genomes. The relative amounts of DNA from nematode and *Wolbachia* therefore remains unaffected after the enrichment step. The preparation of Illumina libraries from Mpe1 and Moz1 samples has been described as part of a previous study (Poole et al. 2019), and a similar protocol was used for Mpe2 and Moz2, using the NEBNext Ultra II FS DNA Library Prep Kit (New England Biolabs Inc., USA) as described by the manufacturer. Following PCR amplification with different index primers to enable multiplexing, the libraries were size selected using NEBNext Sample Purification Beads (NEB cat. # E7767) following manufacturer’s instructions. The approximate insert size and concentration of each library was determined using a 2100 Bioanalyzer with a high sensitivity DNA chip (Agilent Technologies, USA). Two Mpe2 libraries with insert sizes of approximately 500 and 800 bps and two Moz2 libraries with insert sizes of approximately 500 and 950 bps were constructed. Libraries were diluted to 4 nM with 10 mM Tris, 0.1 mM EDTA pH 8 for sequencing. Due to the A:T rich nature of filarial genomes, Phi X DNA was added to balance base pair composition, then sequenced using the Illumina MiSeq and NextSeq500 platforms (paired end, 150 bps).

### Metagenomic assembly and binning

Raw reads were processed using the BBTools package v38.51 (https://sourceforge.net/projects/bbmap/). Duplicate raw reads and bad tiles were removed using the clumpify.sh and filterbytile.sh utilities. Adapter trimming as well as removal of phiX reads and reads with 13-nt or longer stretches of the same nucleotide (poly-G, -C, -A or-T) were performed using bbduk.sh. Human host reads were removed by mapping against the human genome (grch38) using the bbmap.sh utility. The quality metrics of the processed reads at each step were assessed using FastqC v0.11.9 (https://www.bioinformatics.babraham.ac.uk/projects/fastqc/).

Reads from different runs of the libraries prepared from the same genomic DNA sample were combined and used as an input for the assembly of the metagenome using metaSpades v3.14.0 (Nurk et al. 2017). Input reads were mapped back to the assembled scaffolds using bowtie2 v.2.4.1 (Langmead & Salzberg 2012). Binning of metagenomic scaffolds into metagenome assembled genomes of *Mansonella* and *Wolbachia* was performed using the BlobTools v1.1.1 software (Laetsch & Blaxter 2017) and additional manual curation. The bins annotated as “Proteobacteria” and “Nematoda” were analyzed further to retrieve sequences genuinely originating from *Wolbachia*. A cluster of “Proteobacteria” scaffolds in the blobplots of each of the metagenomes was analyzed using blastn against the *w*Cle genome to verify their *Wolbachia* origin. These scaffolds were subsequently collected as respective *Wolbachia* assemblies from each of the isolates. The combined size of scaffolds collected from the Moz1 isolate was only 120 kb, much smaller than a 1 Mb genome typically seen for *Wolbachia,* and so were not analyzed further. Similarly, in the Mpe2 sample, only 12 scaffolds with a total size of 15 kb were identified as candidate *Wolbachia* sequences and were not analyzed further. The tight clusters of “Proteobacteria” scaffolds in the blobplots of Moz2 and Mpe1 were classified as the *w*Moz and *w*Mpe genome assemblies respectively. For the remaining clusters of metagenome scaffolds that were also marked “Proteobacteria” by BlobTools but had a distinctly different %GC as compared to the *Wolbachia* scaffolds, blastn searches against the NCBI-nt database were performed to verify that they did not originate from the *Wolbachia* genome. All the bioinformatics programs were run using the default parameters unless otherwise stated.

The genome sequence of *Wolbachia w*Ocae from the mason bee *Osmia caerulescens* was assembled from Illumina reads available under NCBI SRA accession SRR1221705, originally described in a previous study of arthropod *Wolbachia* (Gerth et al. 2014). The reads were assembled into metagenomic scaffolds using metaSpades. A BlobTools analysis of the resulting assembly was used to identify the cluster of *Wolbachia* scaffolds.

The genome sequence of *Wolbachia w*MoviF from the sheep ked *Melophagus ovinus* was assembled from raw Illumina reads corresponding to the NCBI SRA accession number ERR969522, which originated from a study of sheep ked endosymbionts (Nováková et al. 2015) that were also used recently for *Wolbachia* genome mining (Scholz et al. 2020). In the first round of assembly using all reads as input to metaSpades assembler, potential *Wolbachia* scaffolds were identified using BlobTools. However, a preliminary blastn analysis of these scaffolds indicated a contaminating genome from supergroup A or B *Wolbachia*, in addition to the expected supergroup F *Wolbachia* in this sample. Due to the complicated nature of this dataset, additional iterations of metagenomic assembly and binning were performed using only the reads that mapped to a “bait” database containing all known *Wolbachia* genomes. The mapping of reads to bait databases was performed using bowtie2 (v.2.4.1). The bait database was updated to include the newly assembled sequences, and a new subset of reads was obtained by mapping to this updated database. These reads were then used as input for another round of metagenomic assembly. Candidate *Wolbachia* scaffolds were selected using BlobTools. Whole genome alignments of the assembled scaffolds to three representative and complete reference genomes, namely *w*Cle (supergroup F), *w*Mel (supergroup A), and *w*AlbB (supergroup B) were performed using nucmer v4.0.0beta2 (Marçais et al. 2018, 4) with the parameter “mincluster” set to 100.

### Genome annotation and comparative analysis

For both *w*Mpe and *w*Moz genomes, as well as the *w*Ocae and *w*MoviF genomes, protein-coding genes, rRNA, tRNA, ncRNA genes and pseudogenes were identified using the NCBI prokaryotic genome annotation pipeline (Tatusova et al. 2016). The completeness of the protein-coding content of all 4 genomes was assessed using the BUSCO pipeline v5.1.3 using the “proteobacteria_odb10” reference dataset (Simão et al. 2015). Syntenic blocks between various pairs of genomes were analyzed and visualized using the JupiterPlot tool (https://github.com/JustinChu/JupiterPlot) which uses minimap2 (v 2.17-r941) for whole-genome alignments (Li 2018). The parameters used in Jupiter plots are maxGap = 100,000; minBundleSize = 1,000; m_ref_contig_cutoff = 500; gScaff = 500. The parameter ‘ng’ was set to 100%, so that all the scaffolds from the two assemblies being compared are displayed in the Jupiter plots. Whole-genome alignments of *w*Moz, *w*Mpe, *w*Ocae and *w*MoviF, as well as *w*Mhie, against *w*Cle as the reference genome were performed using the nucmer utility in MUMmer4 with default parameters. The resulting alignment blocks were visualized as concentric Circos plots (Krzywinski et al. 2009) drawn using the R package circlize v0.4.10 (Gu et al. 2014). For global sequence comparisons across multiple *Wolbachia* genomes, the average nucleotide identity (ANI) scores were calculated using the OrthoANIu tool v1.2 (Yoon et al. 2017). The pairwise ANI scores were used for hierarchical clustering of different *Wolbachia,* and a correlation plot was generated using R package corrplot v0.84 (https://github.com/taiyun/corrplot). For a more sensitive measure of sequence similarity and divergence between closely related *Wolbachia*, digital DNA–DNA hybridization (dDDH) scores (Meier-Kolthoff et al. 2013) were computed using the recommended “formula 2” at the “Genome-to-Genome Distance Calculator” (GGDC) web-service (https://ggdc.dsmz.de/ggdc.php). The orthologs of protein coding genes across multiple *Wolbachia* genomes were inferred using OrthoFinder v2.4 (Emms & Kelly 2019).

Insertion sequence (IS) elements were identified via the ISsaga web server (Varani et al. 2011) and by parsing the annotations in GenBank format from NCBI-PGAP pipeline. Other transposons and Group II introns were also inferred via parsing the GenBank files. Functional annotation of protein-coding genes was carried out using the eggNOG-Mapper (Huerta-Cepas et al. 2017) web server (http://eggnogdb.embl.de/#/app/emapper). The analysis of different metabolic pathways was based on the annotations of *w*Cle and *w*Bm genomes available in the KEGG database (Kanehisa et al. 2017).

### Phylogenomic analysis

Phylogenomic analysis was carried out on three distinct sets of input genomes. The first set comprised 127 genomes and included 123 annotated *Wolbachia* genomes available in the NCBI GenBank database (Supplementary Table 1) as well as the 4 genomes reported in this study. For a more robust and refined tree, a second set was used comprising 81 genomes selected from the 127 genomes, excluding incomplete genomes (except those within supergroup F) as well as redundant genomes e.g., multiple isolates of the same *Wolbachia* (Supplementary Table 1). The third set included only the genomes from supergroup F *Wolbachia*, and *w*CfeJ *Wolbachia* from the cat flea *Ctenocephalides felis* as an outgroup.

For each input set, OrthoFinder was used to identify Single Copy Orthologs (SCOs) conserved across all genomes within the set. Protein sequences corresponding to each orthogroup were aligned using mafft v7.149b (Nakamura et al. 2018). The multiple sequence alignments were trimmed using Gblocks v0.91b (Talavera & Castresana 2007). The trimmed blocks from each conserved orthogroup were then concatenated into a supermatrix, and the length of each alignment block was recorded in a nexus format partition file. The corresponding gene sequence tree was generated by collecting the transcript sequences corresponding to the single copy orthologs conserved across all input *Wolbachia*. The DNA sequences corresponding to each conserved orthogroup were first aligned using mafft and then concatenated into a supermatrix sequence, with the lengths of the alignment blocks recorded in a nexus format partition file. Maximum likelihood trees were generated based on protein sequences as well as gene sequences, using the corresponding concatenated supermatrix sequence and partition files (Chernomor et al. 2016) as inputs to iqtree v2.1.2 (Nguyen et al. 2015). Automatic selection of substitution models was performed using ModelFinder (Kalyaanamoorthy et al. 2017) implementation in iqTree (iqTree command line option ‘-m MFP’). Evaluation of branch supports was performed using the iqTree implementations of (i) the SH-like approximate likelihood ratio test (Guindon et al. 2010), and (ii) ultrafast bootstrap (Hoang et al. 2018) with 1000 replicates each (iqTree command-line options ‘-B 1000 -alrt 1000 -abayes -lbp 1000’). The consensus trees were visualized using the iTOL webserver (Letunic & Bork 2021) and further annotated in Adobe Illustrator.

For a phylogenetic analysis of all supergroup F *Wolbachia* based on a combined supermatrix of multiple genes, the sequences for their MLST loci *ftsZ*, *fbpA*, *coxA*, *gatB*, *hcpA* and other loci (*groEL*, *dnaA*, ftsZ and 16S rRNA) were obtained from their genome assemblies and GenBank. The corresponding sequences from *w*CfeJ was used as outgroup. Only the *Wolbachia* with sequences available for at least 3 of these 8 loci were included in this analysis. The multiple sequence alignment of the combined supermatrix of sequences was performed using MUSCLE (Edgar 2004). Phylogenetic tree with automatic model selection, standard bootstrap calculations and single branch tests was generated using iqtree (command-line options -b 1000 -alrt 1000 -abayes).

### Assembly and annotation of *Wolbachia* host genomes and transcriptomes for phylogenomic analysis

The protein coding genes of different *Wolbachia* hosts were obtained as follows. The genomes and proteomes for filarial nematodes *Mansonella perstans*, *Mansonella ozzardi* were recently described (Sinha et al. 2023). Curated proteomes from publicly available data for the filarial hosts *Brugia malayi*, *Brugia pahangi*, *Wuchereria bancrofti*, *Dirofilaria immitis*, *Onchocerca ochengi*, *Onchocerca volvulus*, *Litomosoides sigmodontis*, *Litomosoides brasiliensis*, *Dipetalonema caudispina*, *Cruorifilaria tuberocauda* were the same as recently (Sinha et al. 2023). The genome sequence of *Osmia caerulescens* was obtained from the metagenomic assembly and blobplot analysis for *w*Ocae described above. Annotated genes for *Cimex lectularius* and *Folsomia candida* were obtained from NCBI assemblies GCF_000648675.2 and GCF_002217175.1 respectively. The *Pentalonia nigronervosa* genome was obtained from NCBI accession GCA_014851325.1. Modeling and masking of genomic repeats in *O. caerulescens* and *P. nigronervosa* genomes was performed using RepeatModeler version 2.0.3 (Flynn et al., 2020) and Dfam database (Storer et al., 2021). These repeats were soft-masked in the respective genomes using RepeatMasker 4.1.2-p1 (http://repeatmasker.org/). Gene predictions were carried out on WebAUGUSTUS online server () with the soft-masked genome as input. The soft-masked genomes obtained were used as input to gene prediction using online AUGUSTS web server (Hoff & Stanke 2013). The transcriptomes for *M. ovinus* and *Cimex hemipterus* were assembled *de novo* from SRA datasets SRR17267914 (Zhang et al. 2023) and SRR19217017 respectively. The reads were corrected using rcorrector (Song & Florea 2015) and assembled using Trinity-v2.8.5 (Haas et al. 2013). The longest super-transcripts were selected using transdecoder v5.5.0 (https://github.com/TransDecoder/TransDecoder), and arthropod sequences were selected using kraken version 2.0.9-beta (Wood & Salzberg 2014) and kraken-tools script extract_kraken_reads.py (Lu et al. 2022). Conserved orthologs and species tree for these nematode and arthropod hosts were obtained from OrthoFinder (Emms & Kelly 2019).

### Gene loss and pseudogene analysis

OrthoFinder results, from the set of 81 genomes described above, were analyzed to identify gene loss events across different *Wolbachia*, by cataloging orthogroups which had gene members present in all arthropod *Wolbachia* but absent in filarial *Wolbachia*.

Pseudogenization of protein-coding genes is a major factor contributing to gene loss. Pseudogenes common across different supergroup F *Wolbachia* were identified as follows. A catalog of pseudogenes for each *Wolbachia* was first obtained from its NCBI-PGAP annotation. Next, for each pseudogene, the GenBank accession for its most likely intact ancestral protein was also obtained from the NCBI-PGAP annotations. For a phylogenetically consistent and robust analysis, instead of comparing these protein accessions directly, the corresponding ancestral orthogroup was retrieved from the results of OrthoFinder and these “pseudogene-orthogroups” were compared across different *Wolbachia*. If the same orthogroup was found to be pseudogenized in more than one *Wolbachia*, the corresponding pseudogenes were designated as shared or common pseudogenes. The overlap between the ancestral pseudogene-orthogroups across different *Wolbachia* was visualized using the R package UpsetR (Conway et al. 2017). Pseudogenes derived from transposons and IS elements were excluded from this analysis.

The dN/dS values for *w*Bpa-bfr and *w*Mhie-*bfr* loci were calculated from pairwise alignments with intact genes from closely related *Wolbachia*, using PAL2NAL utility (Suyama et al. 2006). The substitution saturation index of both these loci was evaluated using Xia’s test (Xia et al. 2003) using DAMBE (Xia 2018). The proportion of invariant sites at *bfr* locus for this analysis was obtained via PhyML (Guindon et al. 2010) analysis of 13 intact bacterioferritin genes from all supergroups. A global analysis of dN/dS values for all genes and pseudogenes for *w*Bpa and *w*Bma against the *w*ChemPL13 reference genome were obtained using Pseudofinder (Syberg-Olsen et al. 2022).

### Annotation of phage derived elements and cytoplasmic incompatibility factors

Phage-derived genes and pseudogenes were annotated following the latest taxonomic classification scheme of *Wolbachia* phages (Bordenstein & Bordenstein 2022). The sequences of *Wolbachia* phage genes and pseudogenes were obtained from the Phage-WO database available at https://lab.vanderbilt.edu/bordenstein/phage-wo/database. Their best sequence matches in the proteomes of the set of 81 *Wolbachia* genomes were first identified using blastp and blastx searches. The orthogroups corresponding to these best hits were considered as phage-derived orthogroups, and the corresponding orthologs in supergroup F *Wolbachia* were annotated as phage-derived genes. Genes encoding IS-elements were excluded from this analysis to avoid mismatches and overcounting arising from the repeated and fragmented nature of genes in these families. Pseudogenes were annotated as phage-derived if their predicted ancestral proteins had an ortholog in the phage WO database. For *cifA* and *cifB* gene annotations, reference gene sequences were compiled from published data (Martinez et al. 2021) and used in blast searches against the proteomes from the 81 *Wolbachia* genomes used in the phylogenomic analysis above, and the corresponding orthogroups and orthologs within supergroup F *Wolbachia* were annotated as *cifA* and *cifB*.

## Acknowledgments

The authors thank Jeremy Foster and Sofia Roitman for helpful discussions and comments on the manuscript, and Laurie Mazzola and Danielle Fuchs from the DNA sequencing core at New England Biolabs. The work was inspired by Don Comb and funded by New England Biolabs. Field research in Brazil was supported by the Fundação de Amparo à Pesquisa do Estado de São Paulo (FAPESP), research grant to M.U.F. (2013/12723-7) and doctoral scholarship to N.F.L. (2013/ 26928-0).

## Author Contributions

N.F. L., M. U. F., F.F. F., S.W. collected the clinical samples. A.S., Z. L, C.B. P., L. E. performed the sequencing experiments and bioinformatics analysis. A.S., Z. L, C.B. P., L.E., C.K.S.C. analyzed the data. A.S., Z. L, C.B. P., C.K.S.C wrote the initial manuscript. A.S., Z. L, C.B. P., L.E., M. U. F., S.W., C.K.S.C and all authors contributed to the final manuscript.

## Data Availability Statement

Raw Illumina reads for the *M. perstans* isolate Mpe1 and the *M. ozzardi* isolate Moz2 have been deposited in the NCBI SRA database under BioProject accession numbers PRJNA666672 and PRJNA666671 respectively. The assembled genome and PGAP annotations for the *Wolbachia w*Mpe and *w*Moz are available from the NCBI GenBank database with accession numbers JACZHU000000000 and JADAKK000000000 respectively. The annotated genomes for *w*Ocae and *w*MoviF are available as GenBank accessions JAHDJT000000000 and JAHRBA000000000 respectively, deposited under BioProject accessions PRJNA244005 and PRJNA739984 respectively.

## Notes

### Competing Interest Statement

The authors have declared no competing interest.

### Summary of Updates

1. Added co-phylogeny of Wolbachia and their filarial and arthropod hosts (revised Figure 5). 2. Added dN/dS calculations for bacterioferritin pseudogene in wBpa. 3. Supplemental files updated.

