## Supplementary Figure for "Multiple origins of nematode-*Wolbachia* symbiosis in supergroup F and convergent loss of bacterioferritin in filarial *Wolbachia*"

### Slide 1
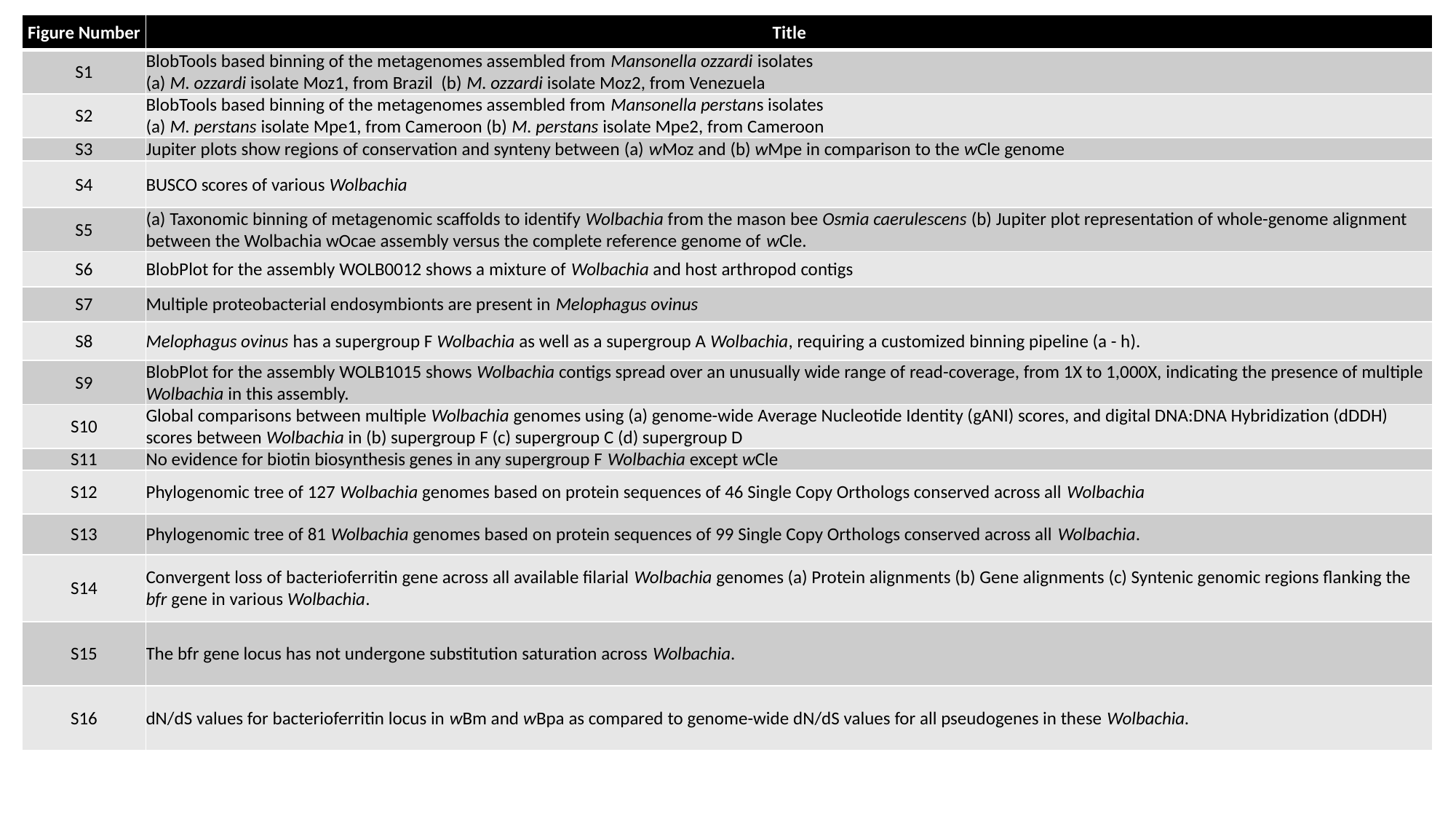

| Figure Number | Title |
| --- | --- |
| S1 | BlobTools based binning of the metagenomes assembled from Mansonella ozzardi isolates (a) M. ozzardi isolate Moz1, from Brazil (b) M. ozzardi isolate Moz2, from Venezuela |
| S2 | BlobTools based binning of the metagenomes assembled from Mansonella perstans isolates (a) M. perstans isolate Mpe1, from Cameroon (b) M. perstans isolate Mpe2, from Cameroon |
| S3 | Jupiter plots show regions of conservation and synteny between (a) wMoz and (b) wMpe in comparison to the wCle genome |
| S4 | BUSCO scores of various Wolbachia |
| S5 | (a) Taxonomic binning of metagenomic scaffolds to identify Wolbachia from the mason bee Osmia caerulescens (b) Jupiter plot representation of whole-genome alignment between the Wolbachia wOcae assembly versus the complete reference genome of wCle. |
| S6 | BlobPlot for the assembly WOLB0012 shows a mixture of Wolbachia and host arthropod contigs |
| S7 | Multiple proteobacterial endosymbionts are present in Melophagus ovinus |
| S8 | Melophagus ovinus has a supergroup F Wolbachia as well as a supergroup A Wolbachia, requiring a customized binning pipeline (a - h). |
| S9 | BlobPlot for the assembly WOLB1015 shows Wolbachia contigs spread over an unusually wide range of read-coverage, from 1X to 1,000X, indicating the presence of multiple Wolbachia in this assembly. |
| S10 | Global comparisons between multiple Wolbachia genomes using (a) genome-wide Average Nucleotide Identity (gANI) scores, and digital DNA:DNA Hybridization (dDDH) scores between Wolbachia in (b) supergroup F (c) supergroup C (d) supergroup D |
| S11 | No evidence for biotin biosynthesis genes in any supergroup F Wolbachia except wCle |
| S12 | Phylogenomic tree of 127 Wolbachia genomes based on protein sequences of 46 Single Copy Orthologs conserved across all Wolbachia |
| S13 | Phylogenomic tree of 81 Wolbachia genomes based on protein sequences of 99 Single Copy Orthologs conserved across all Wolbachia. |
| S14 | Convergent loss of bacterioferritin gene across all available filarial Wolbachia genomes (a) Protein alignments (b) Gene alignments (c) Syntenic genomic regions flanking the bfr gene in various Wolbachia. |
| S15 | The bfr gene locus has not undergone substitution saturation across Wolbachia. |
| S16 | dN/dS values for bacterioferritin locus in wBm and wBpa as compared to genome-wide dN/dS values for all pseudogenes in these Wolbachia. |

### Slide 2
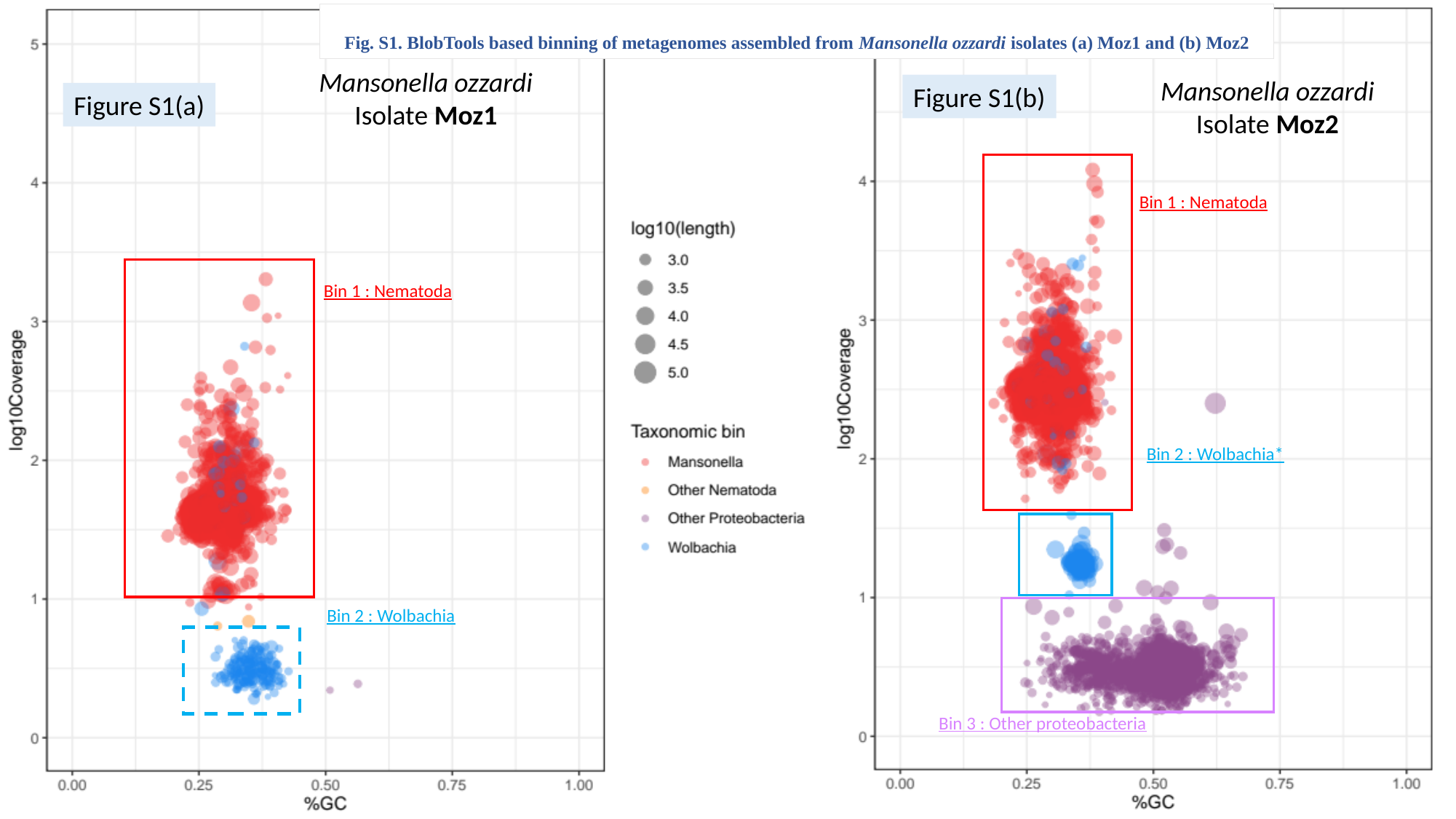

Figure S1(b)
Figure S1(a)
Fig. S1. BlobTools based binning of metagenomes assembled from Mansonella ozzardi isolates (a) Moz1 and (b) Moz2
Mansonella ozzardi
Isolate Moz1
Bin 1 : Nematoda
Bin 2 : Wolbachia
Mansonella ozzardi
Isolate Moz2
Bin 1 : Nematoda
Bin 2 : Wolbachia*
Bin 3 : Other proteobacteria

### Slide 3
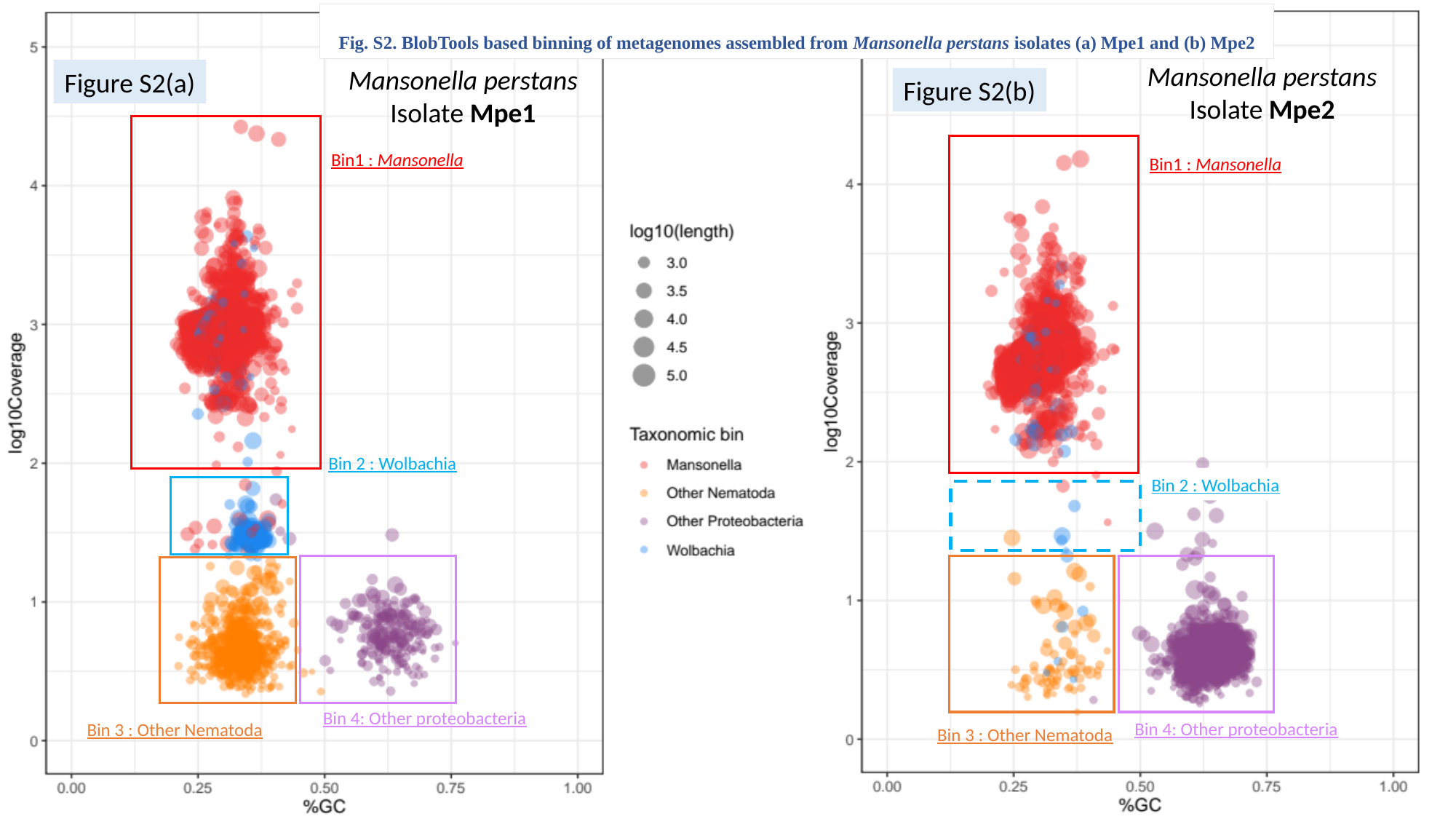

Fig. S2. BlobTools based binning of metagenomes assembled from Mansonella perstans isolates (a) Mpe1 and (b) Mpe2
Mansonella perstans
Isolate Mpe1
Bin1 : Mansonella
Bin 2 : Wolbachia
Bin 4: Other proteobacteria
Bin 3 : Other Nematoda
Mansonella perstans
Isolate Mpe2
Bin1 : Mansonella
Bin 2 : Wolbachia
Bin 4: Other proteobacteria
Bin 3 : Other Nematoda
Figure S2(a)
Figure S2(b)

### Slide 4
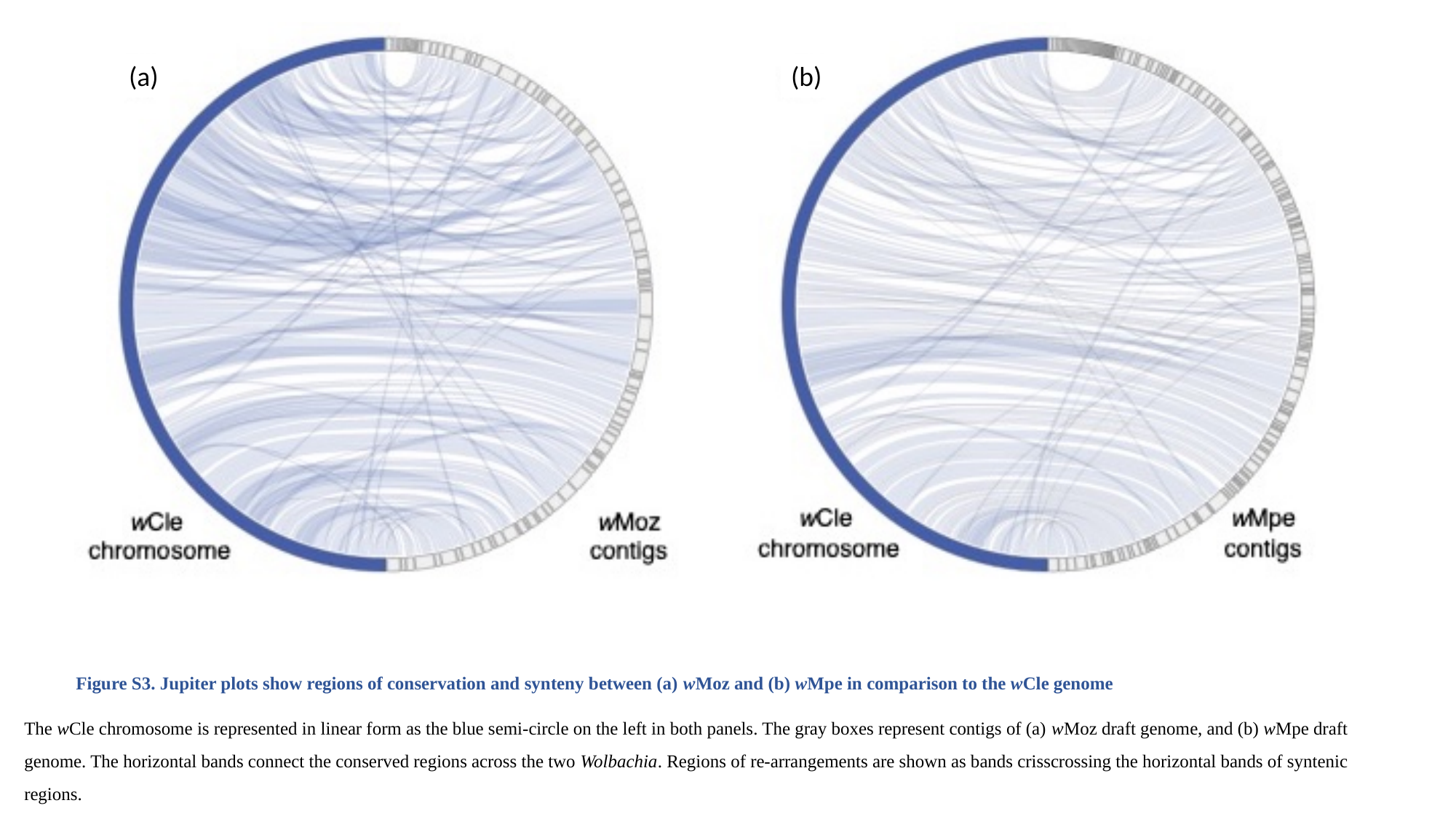

(a)
(b)
Figure S3. Jupiter plots show regions of conservation and synteny between (a) wMoz and (b) wMpe in comparison to the wCle genome​
The wCle chromosome is represented in linear form as the blue semi-circle on the left in both panels. The gray boxes represent contigs of (a) wMoz draft genome, and (b) wMpe draft genome. The horizontal bands connect the conserved regions across the two Wolbachia. Regions of re-arrangements are shown as bands crisscrossing the horizontal bands of syntenic regions. ​

### Slide 5
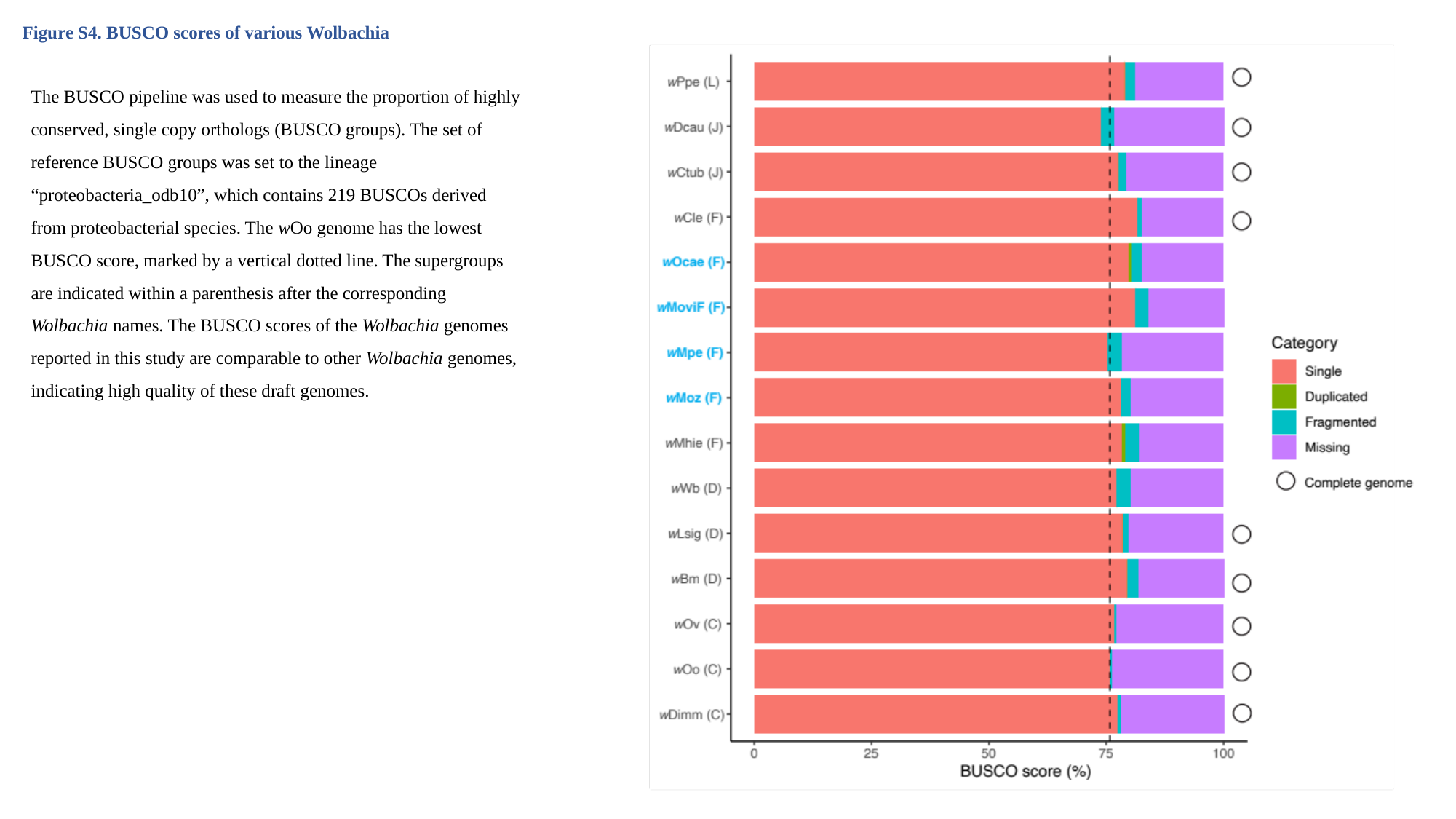

Figure S4. BUSCO scores of various Wolbachia
The BUSCO pipeline was used to measure the proportion of highly conserved, single copy orthologs (BUSCO groups). The set of reference BUSCO groups was set to the lineage “proteobacteria_odb10”, which contains 219 BUSCOs derived from proteobacterial species. The wOo genome has the lowest BUSCO score, marked by a vertical dotted line. The supergroups are indicated within a parenthesis after the corresponding Wolbachia names. The BUSCO scores of the Wolbachia genomes reported in this study are comparable to other Wolbachia genomes, indicating high quality of these draft genomes.

### Slide 6
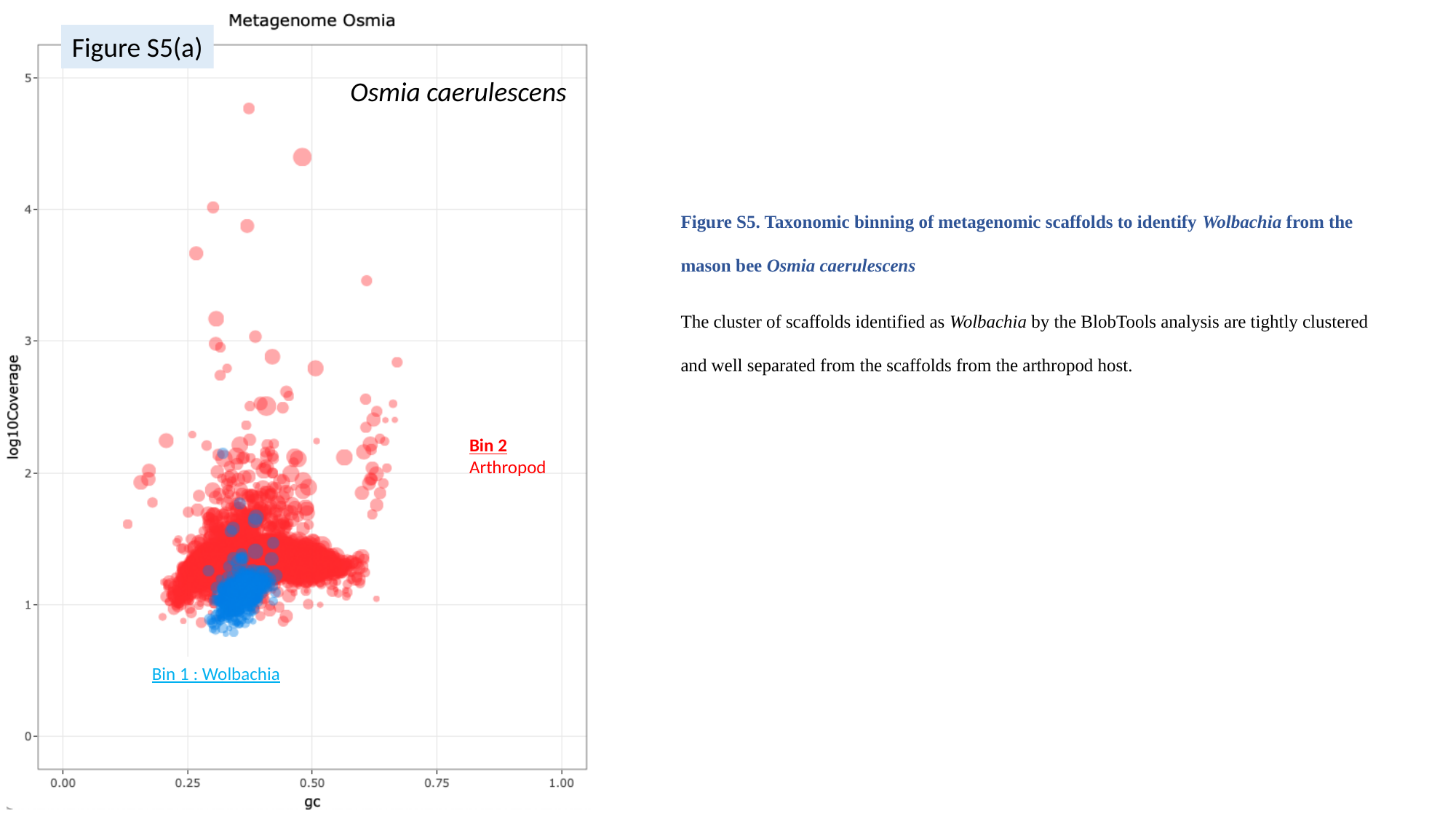

Figure S5(a)
Osmia caerulescens
Figure S5. Taxonomic binning of metagenomic scaffolds to identify Wolbachia from the mason bee Osmia caerulescens
The cluster of scaffolds identified as Wolbachia by the BlobTools analysis are tightly clustered and well separated from the scaffolds from the arthropod host.
Bin 2
Arthropod
Bin 1 : Wolbachia

### Slide 7
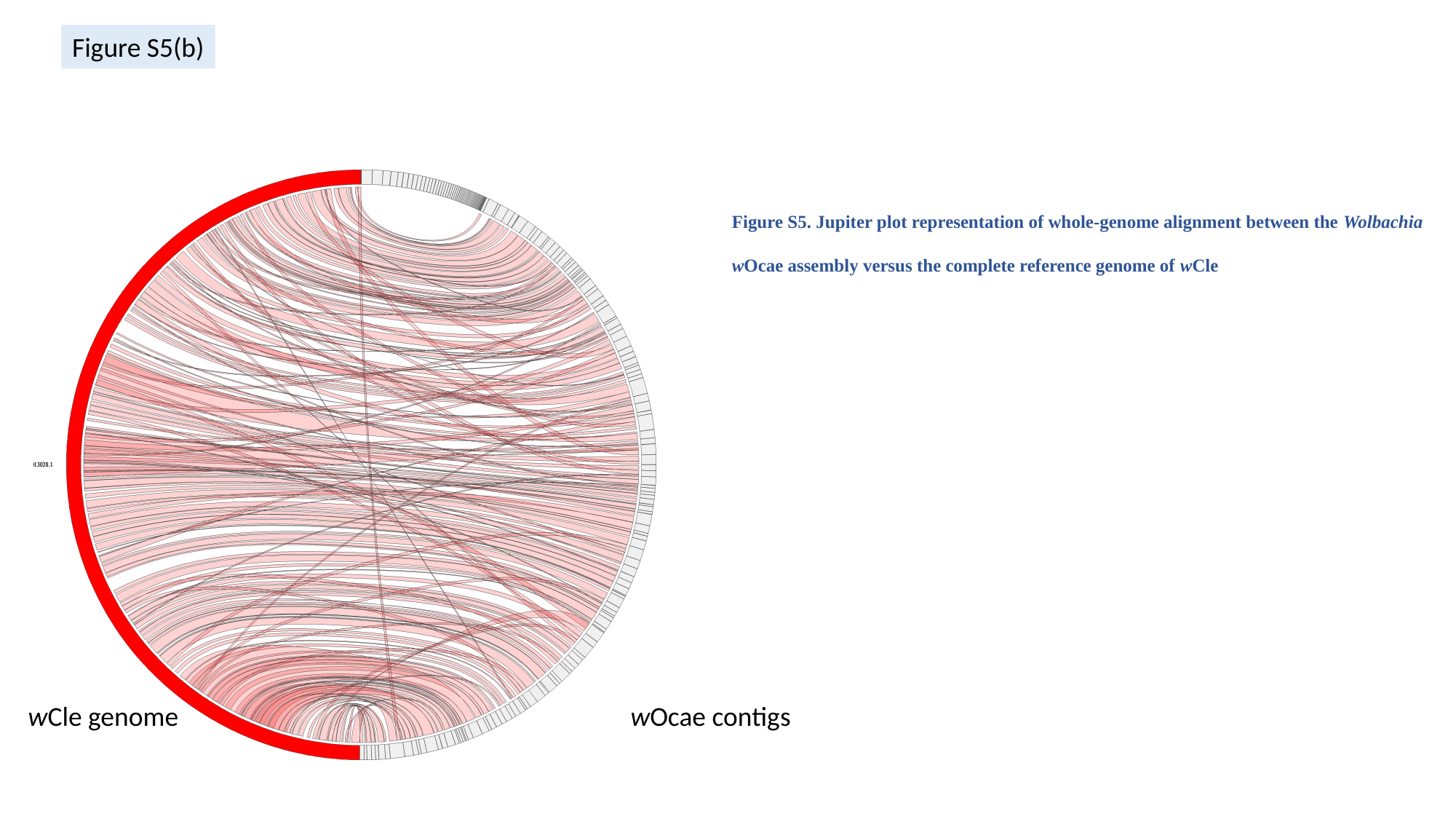

Figure S5(b)
Figure S5. Jupiter plot representation of whole-genome alignment between the Wolbachia wOcae assembly versus the complete reference genome of wCle
wCle genome
wOcae contigs

### Slide 8
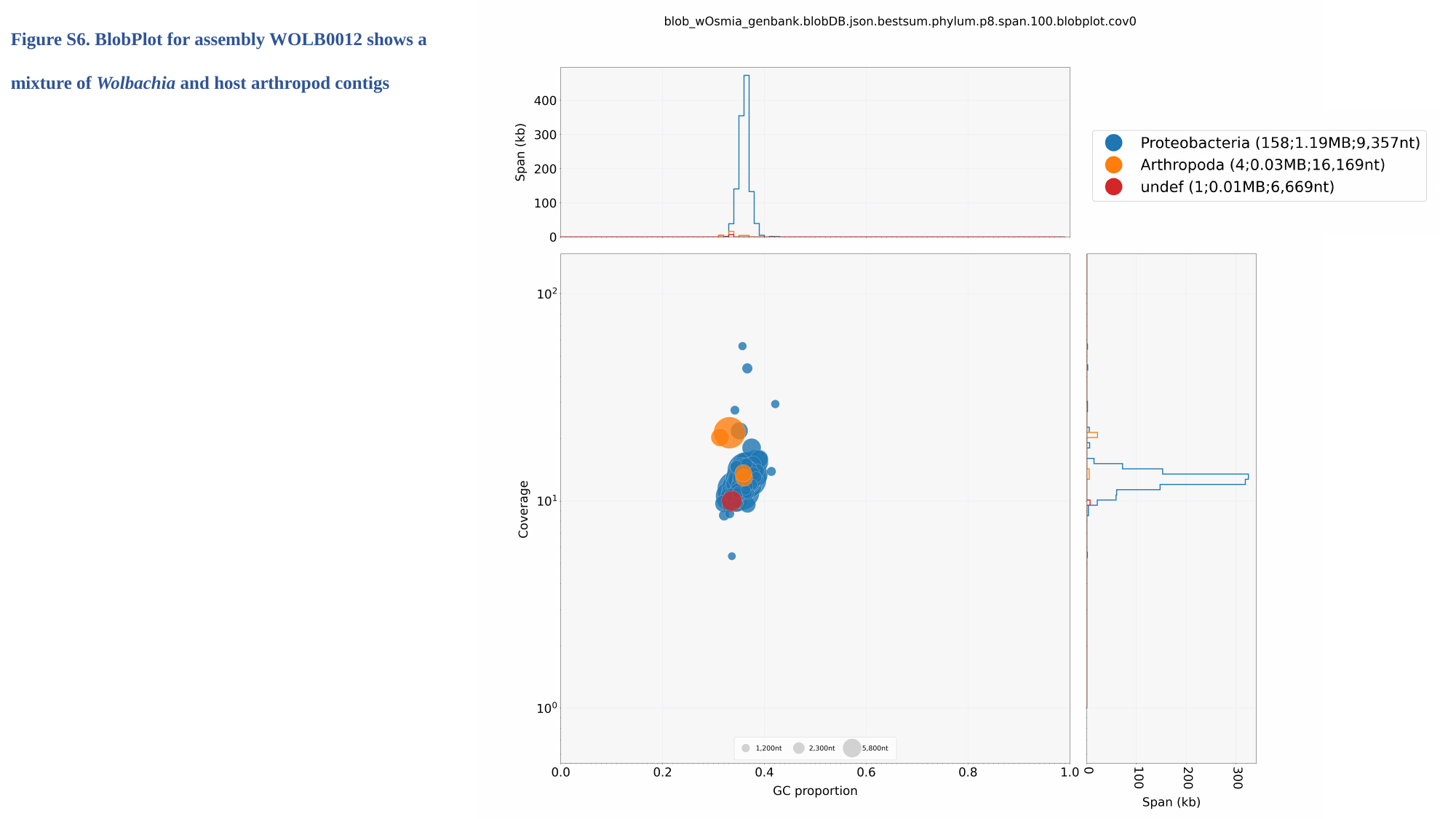

Figure S6. BlobPlot for assembly WOLB0012 shows a mixture of Wolbachia and host arthropod contigs

### Slide 9
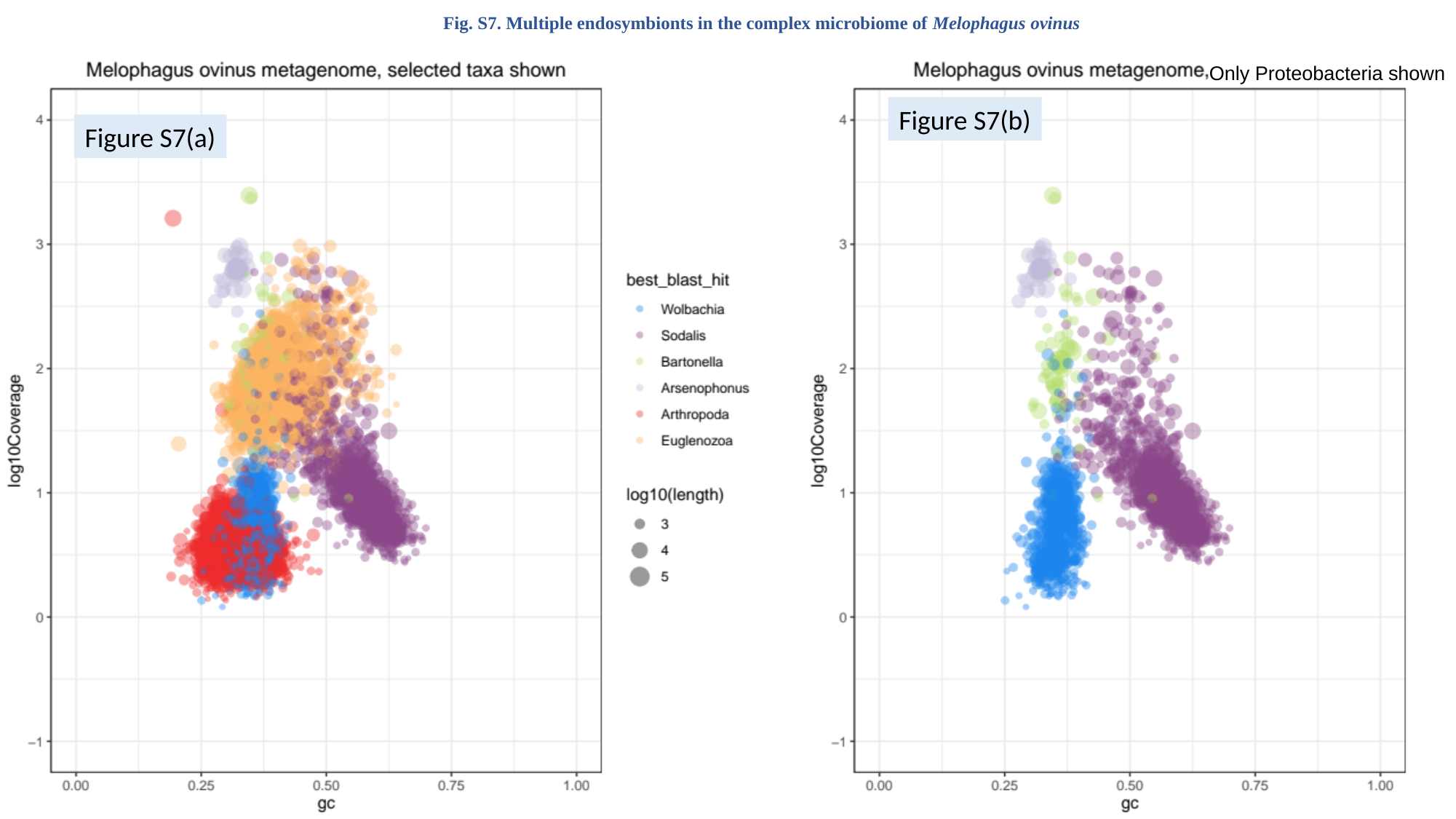

Fig. S7. Multiple endosymbionts in the complex microbiome of Melophagus ovinus
Only Proteobacteria shown
Figure S7(b)
Figure S7(a)

### Slide 10
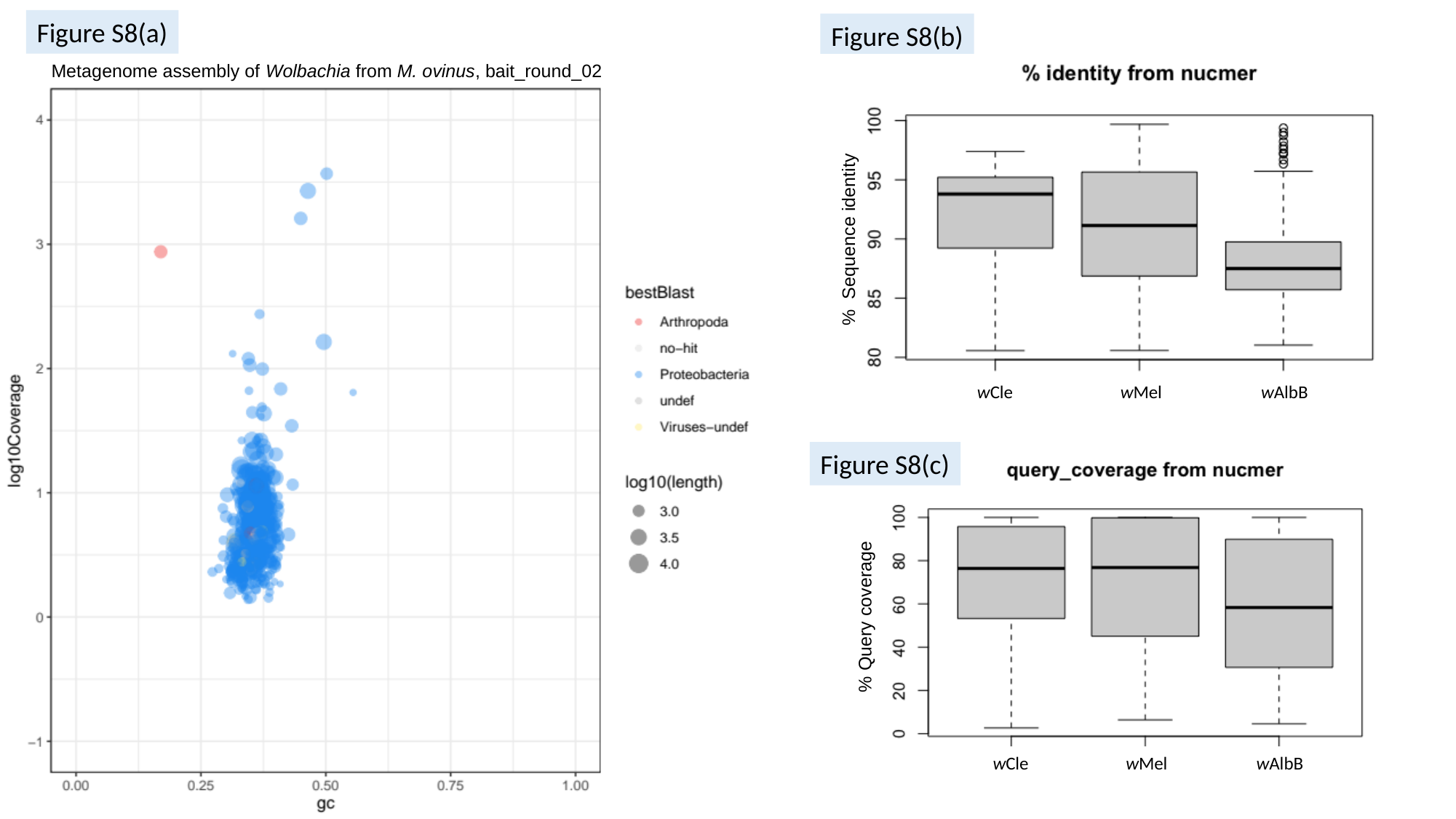

Figure S8(a)
Figure S8(b)
Metagenome assembly of Wolbachia from M. ovinus, bait_round_02
% Sequence identity
wCle
wMel
wAlbB
Figure S8(c)
% Query coverage
wCle
wMel
wAlbB

### Slide 11
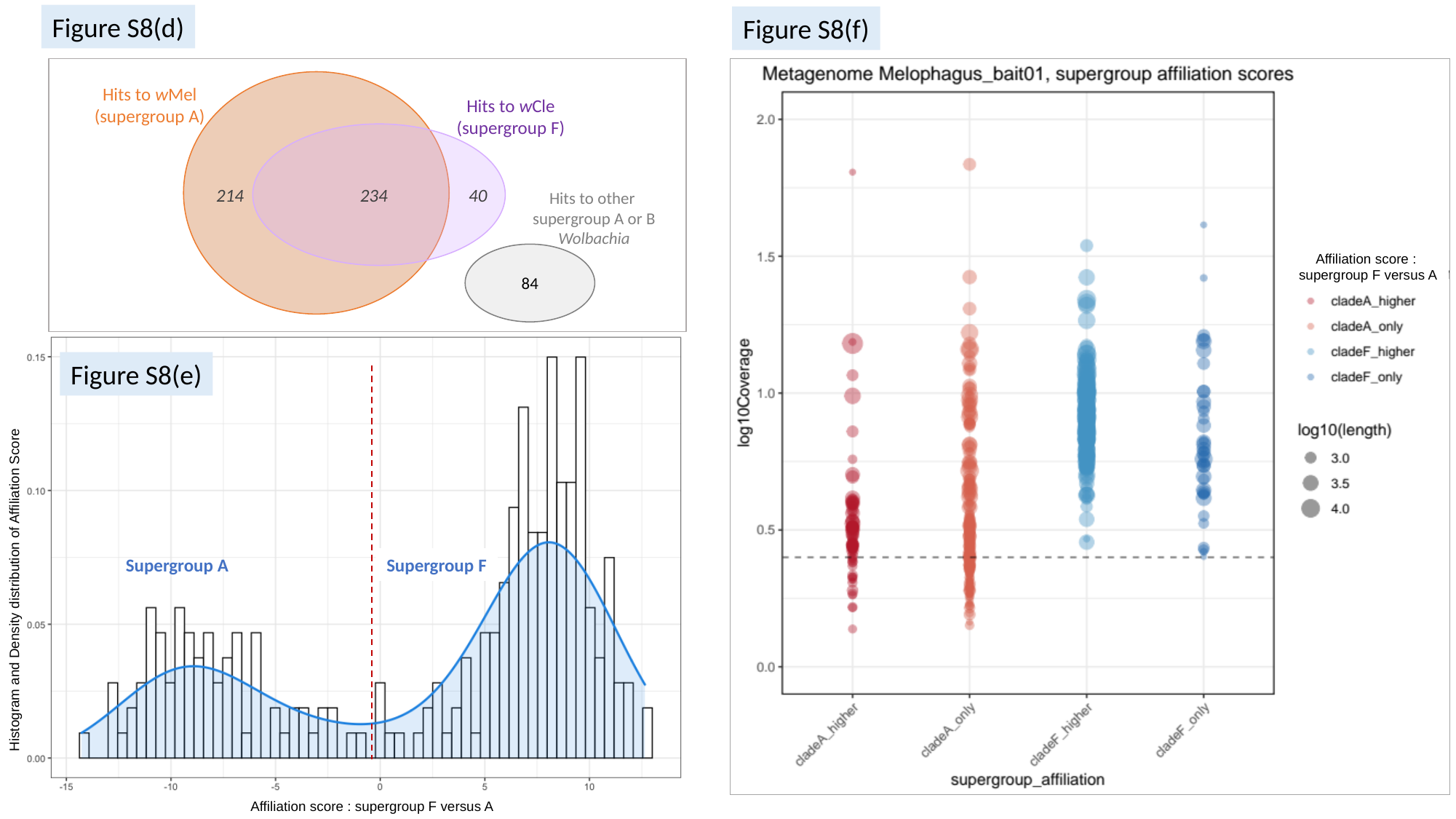

Figure S8(d)
Figure S8(f)
Hits to wMel
(supergroup A)
Hits to wCle
(supergroup F)
214
234
40
Affiliation score :
supergroup F versus A
Hits to other supergroup A or B Wolbachia
84
Supergroup A
Supergroup F
Histogram and Density distribution of Affiliation Score
Affiliation score : supergroup F versus A
Figure S8(e)

### Slide 12
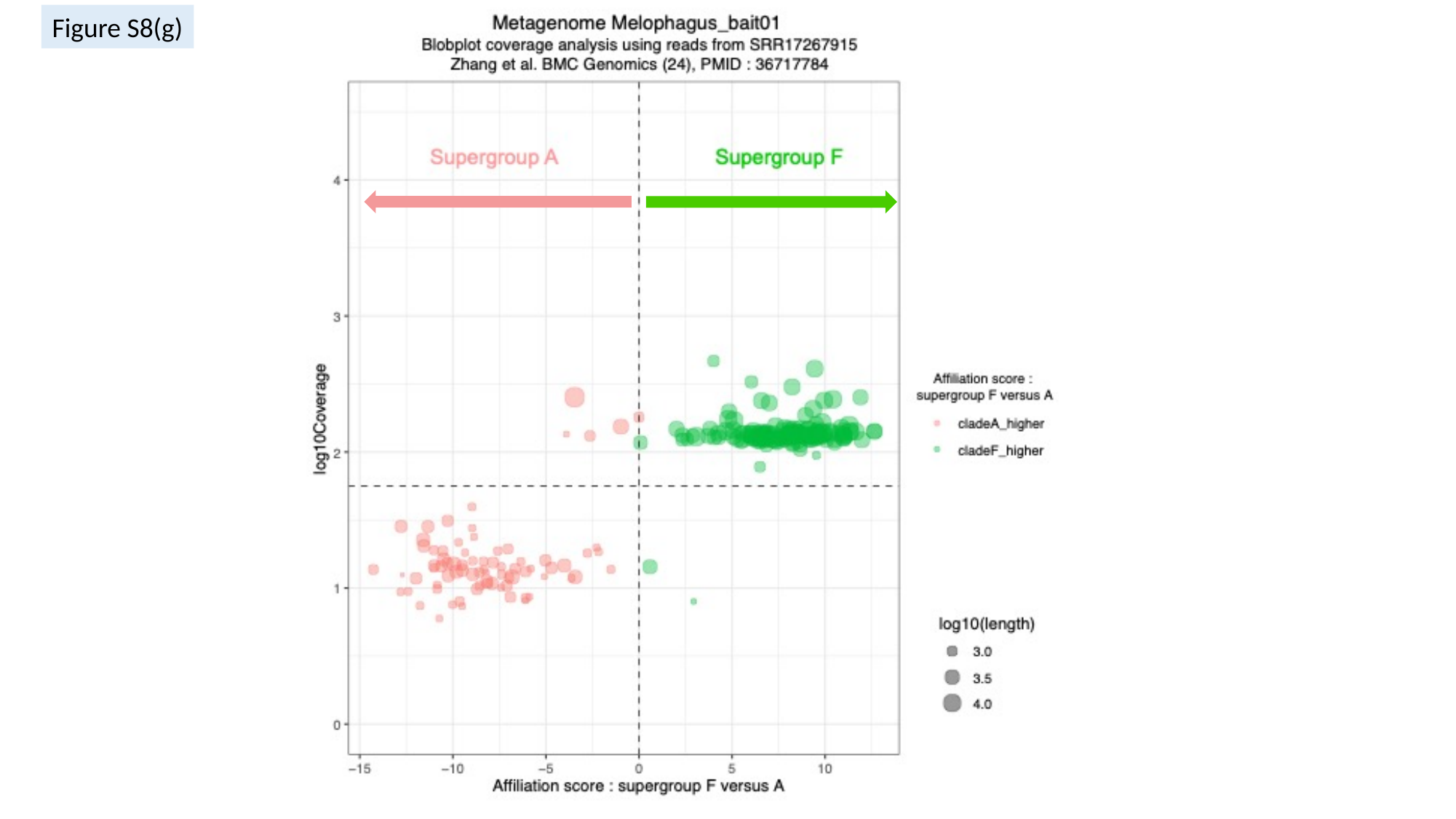

Figure S8(g)

### Slide 13
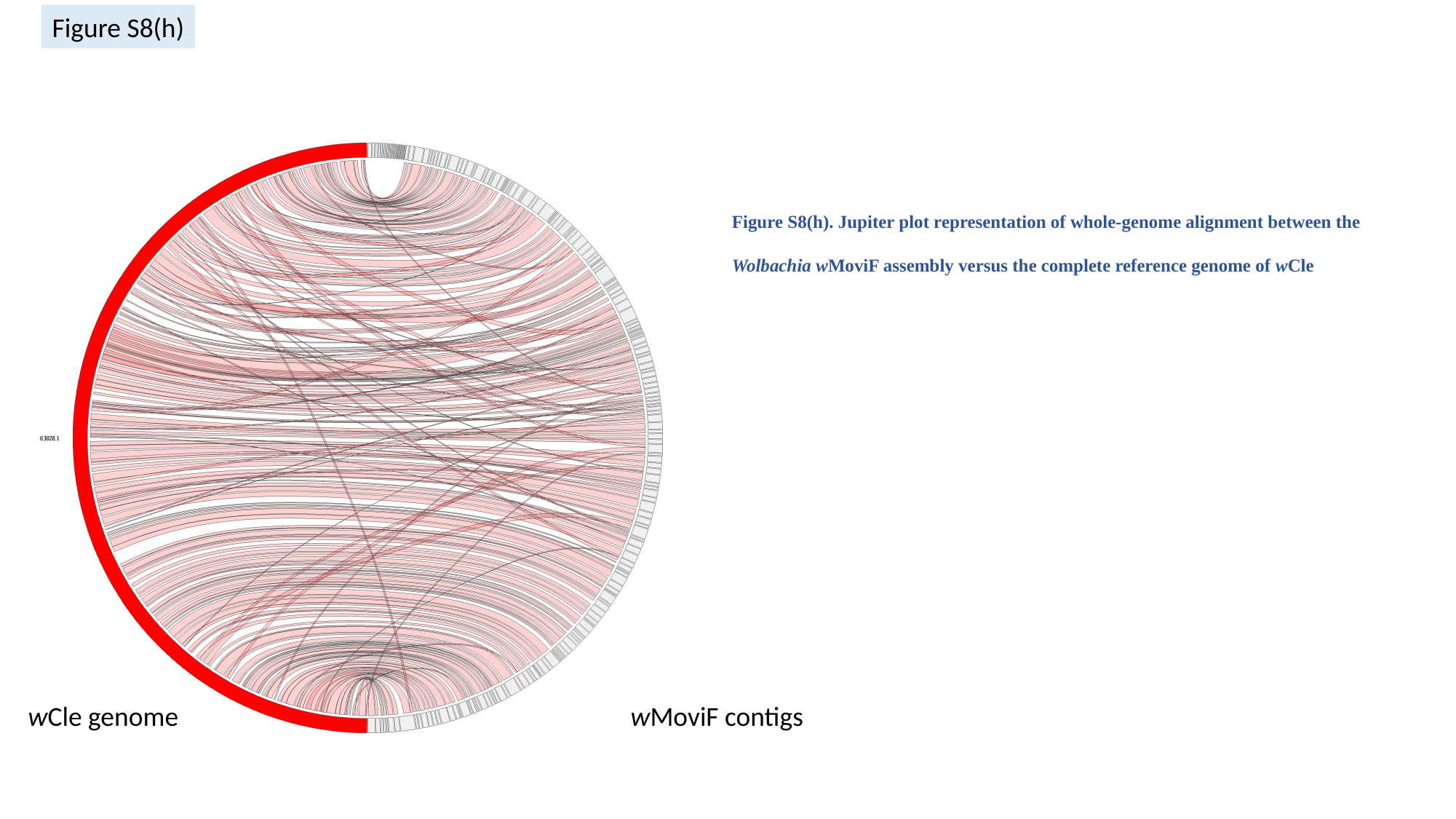

Figure S8(h)
Figure S8(h). Jupiter plot representation of whole-genome alignment between the Wolbachia wMoviF assembly versus the complete reference genome of wCle
wCle genome
wMoviF contigs

### Slide 14
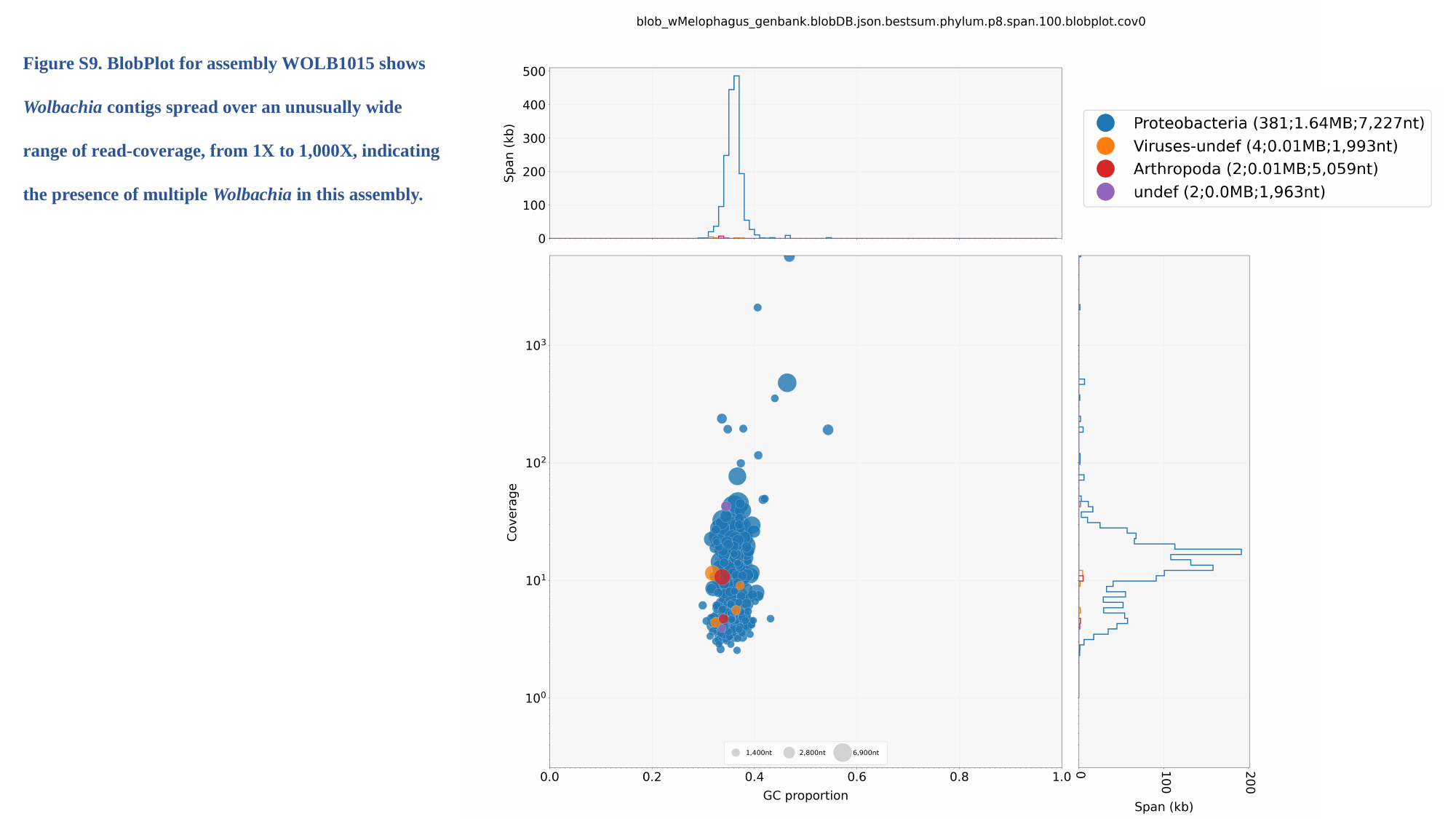

Figure S9. BlobPlot for assembly WOLB1015 shows Wolbachia contigs spread over an unusually wide range of read-coverage, from 1X to 1,000X, indicating the presence of multiple Wolbachia in this assembly.

### Slide 15
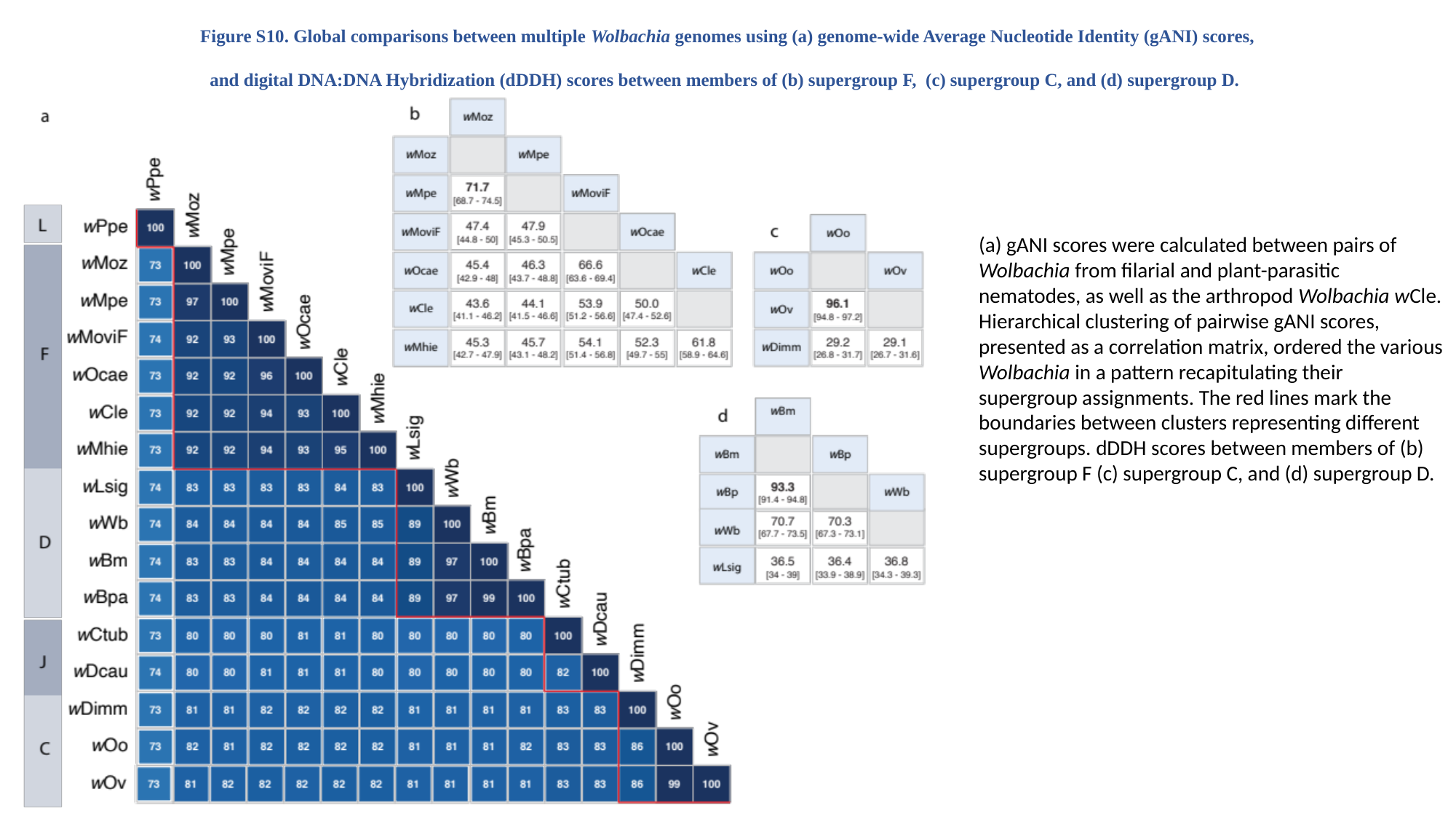

Figure S10. Global comparisons between multiple Wolbachia genomes using (a) genome-wide Average Nucleotide Identity (gANI) scores, and digital DNA:DNA Hybridization (dDDH) scores between members of (b) supergroup F, (c) supergroup C, and (d) supergroup D.
(a) gANI scores were calculated between pairs of Wolbachia from filarial and plant-parasitic nematodes, as well as the arthropod Wolbachia wCle. Hierarchical clustering of pairwise gANI scores, presented as a correlation matrix, ordered the various Wolbachia in a pattern recapitulating their supergroup assignments. The red lines mark the boundaries between clusters representing different supergroups. dDDH scores between members of (b) supergroup F (c) supergroup C, and (d) supergroup D.

### Slide 16
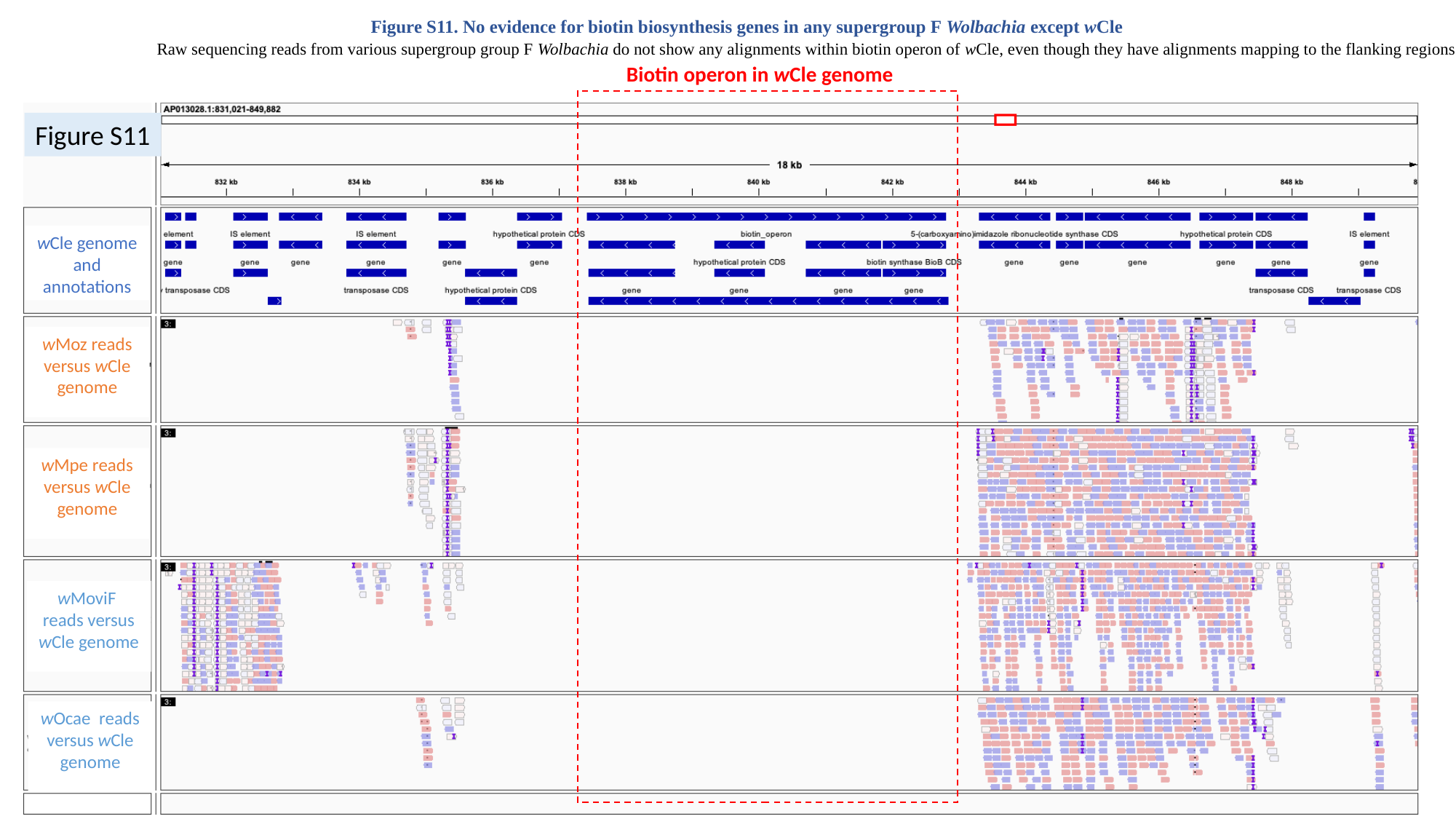

Figure S11. No evidence for biotin biosynthesis genes in any supergroup F Wolbachia except wCle
Raw sequencing reads from various supergroup group F Wolbachia do not show any alignments within biotin operon of wCle, even though they have alignments mapping to the flanking regions.
Biotin operon in wCle genome
wCle genome
and annotations
wMoz reads versus wCle genome
wMpe reads versus wCle genome
wMoviF reads versus wCle genome
wOcae reads versus wCle genome
Figure S11

### Slide 17
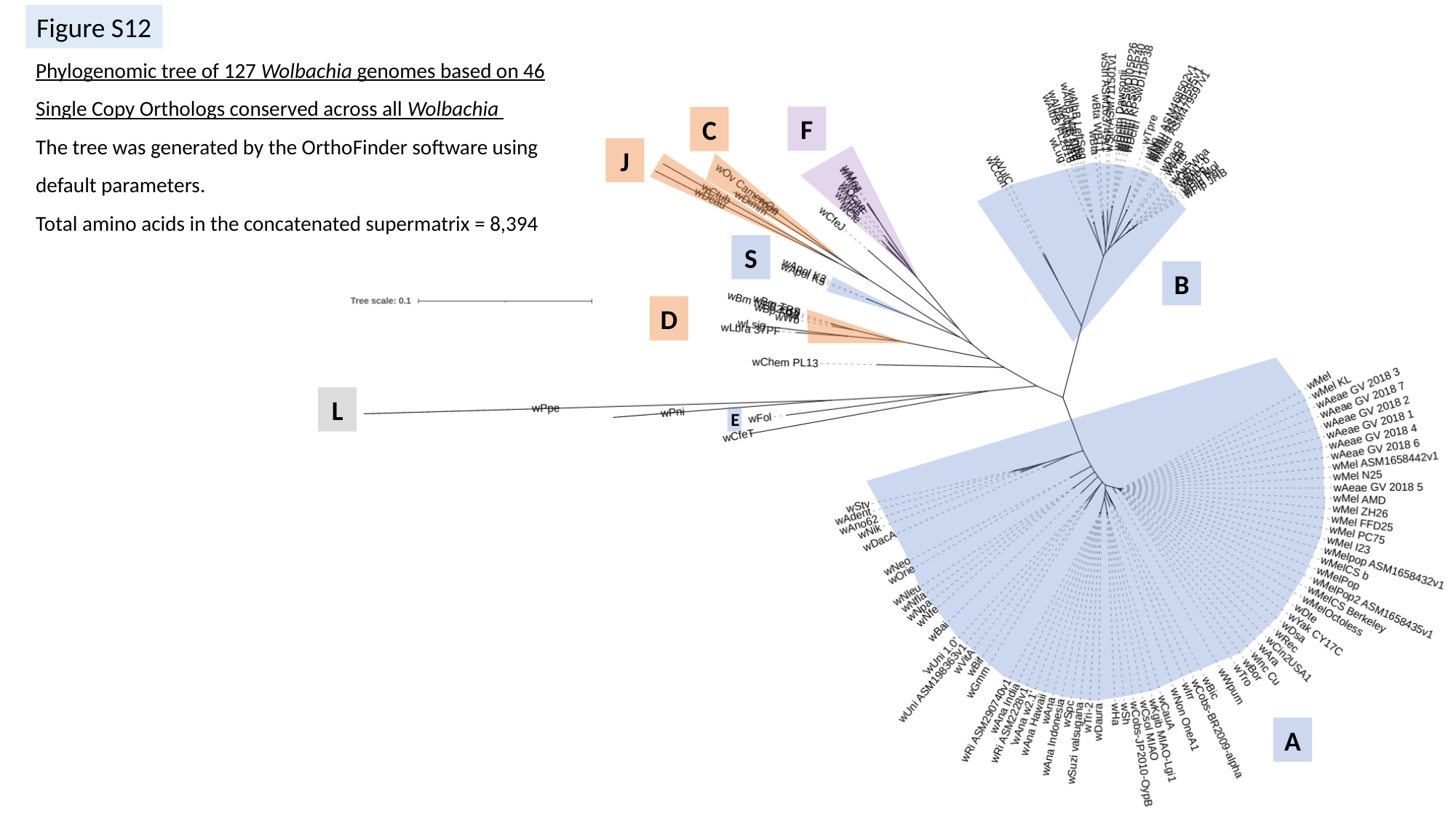

Figure S12
F
C
J
S
B
D
L
A
Phylogenomic tree of 127 Wolbachia genomes based on 46 Single Copy Orthologs conserved across all Wolbachia
The tree was generated by the OrthoFinder software using default parameters.
Total amino acids in the concatenated supermatrix = 8,394
E

### Slide 18
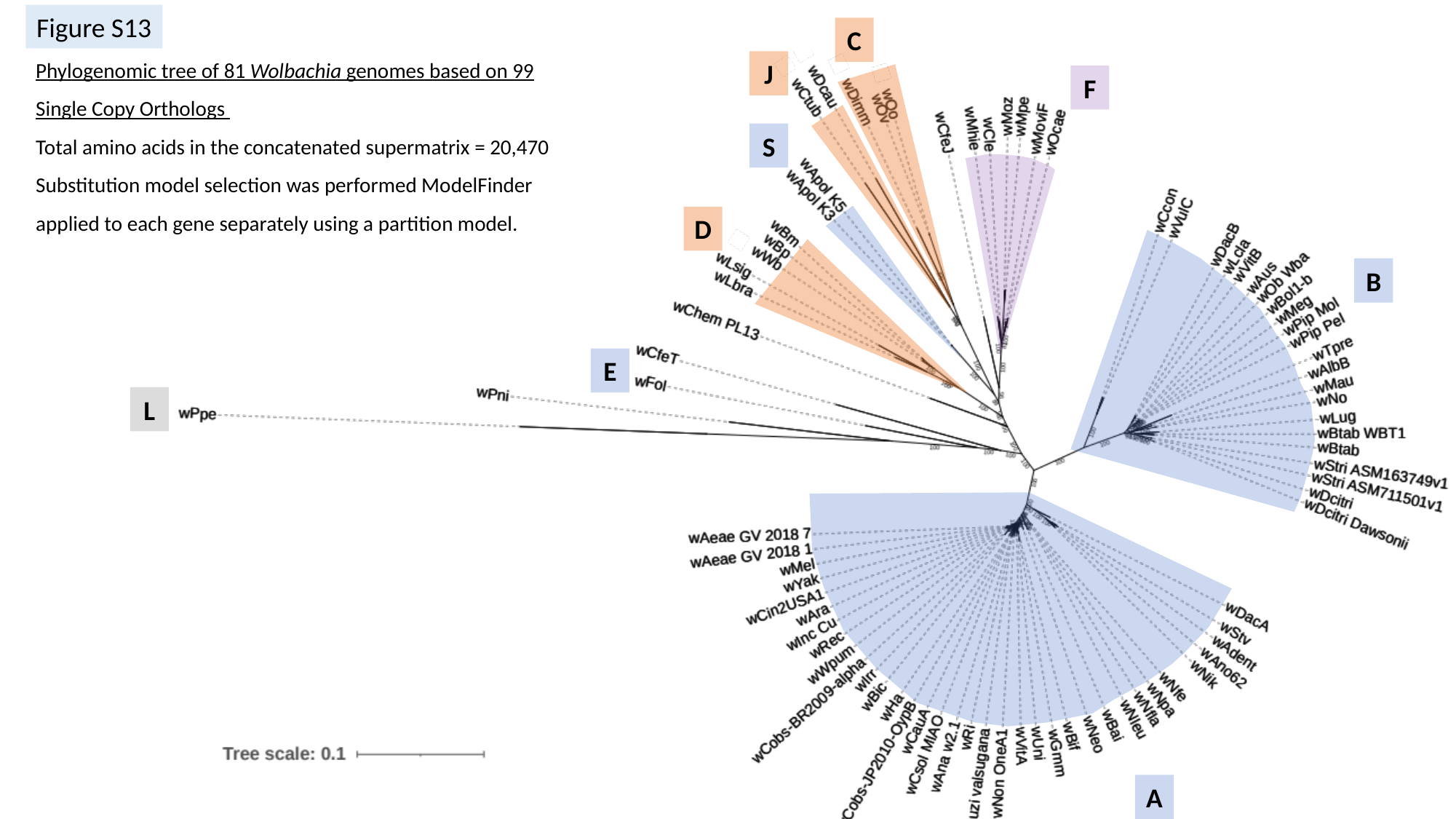

Figure S13
C
J
F
S
D
B
E
L
A
Phylogenomic tree of 81 Wolbachia genomes based on 99 Single Copy Orthologs
Total amino acids in the concatenated supermatrix = 20,470
Substitution model selection was performed ModelFinder applied to each gene separately using a partition model.

### Slide 19
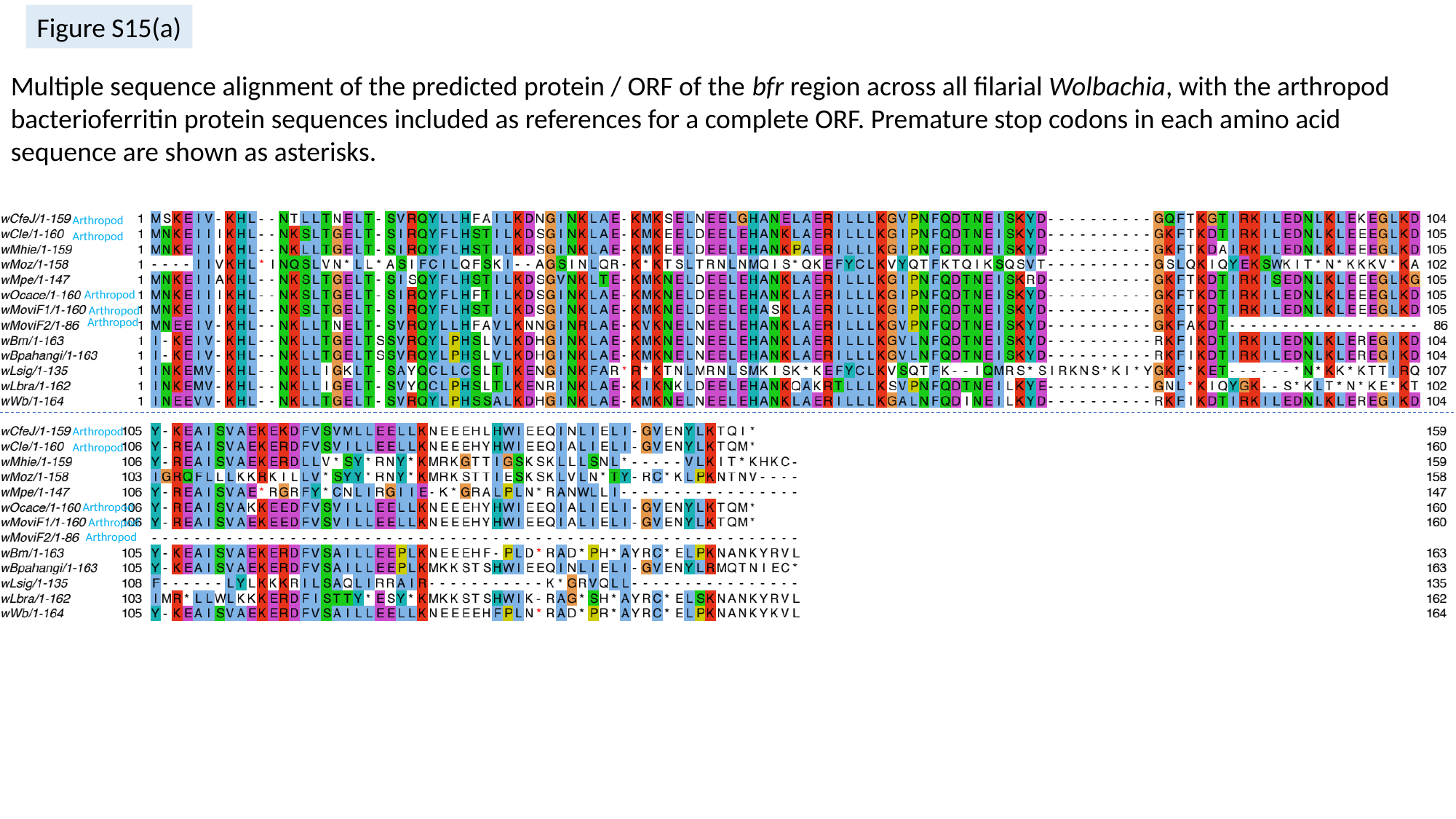

Figure S15(a)
Multiple sequence alignment of the predicted protein / ORF of the bfr region across all filarial Wolbachia, with the arthropod bacterioferritin protein sequences included as references for a complete ORF. Premature stop codons in each amino acid sequence are shown as asterisks.
Arthropod
Arthropod
Arthropod
Arthropod
Arthropod
Arthropod
Arthropod
Arthropod
Arthropod
Arthropod

### Slide 20
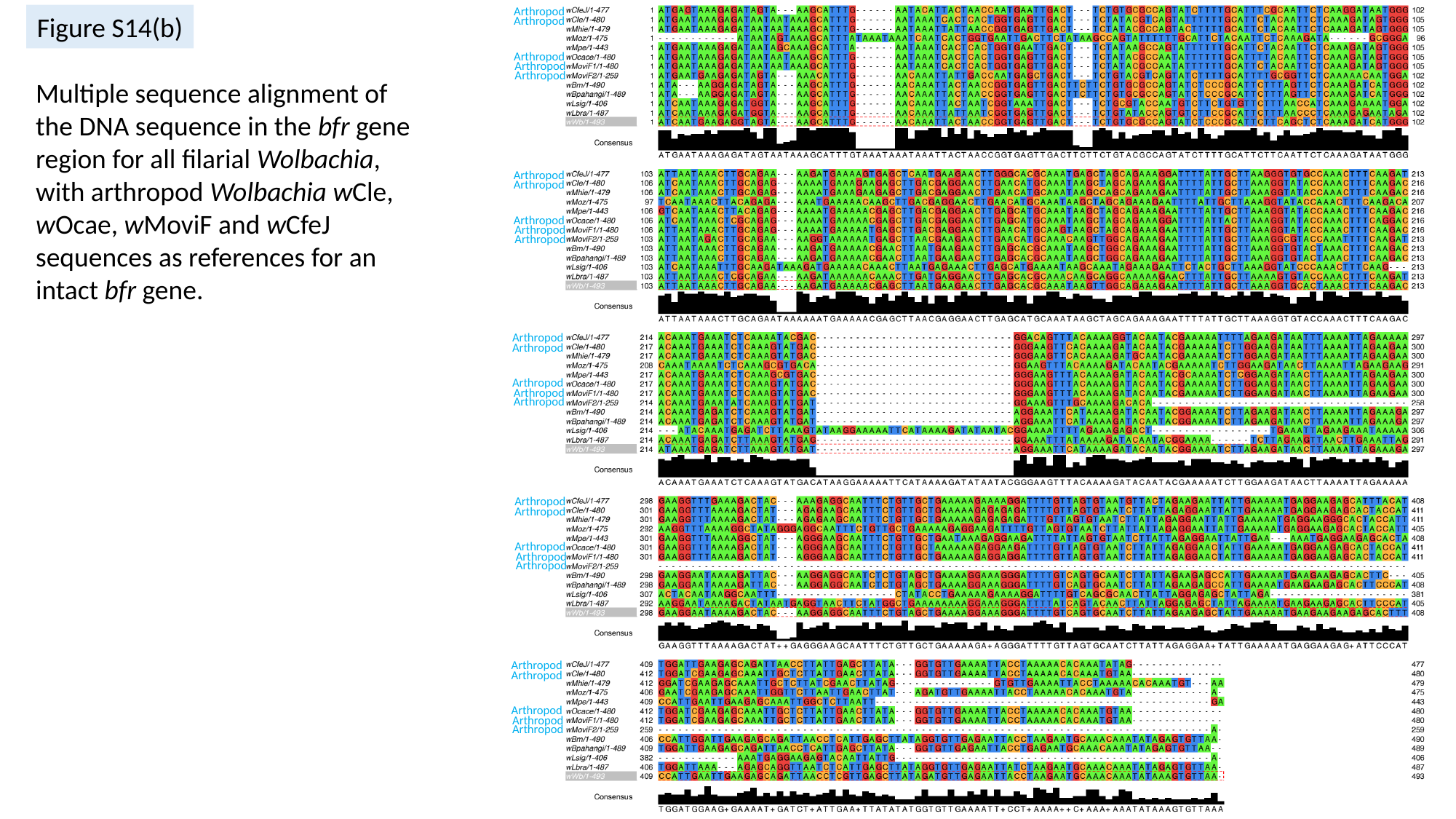

Arthropod
Arthropod
Arthropod
Arthropod
Arthropod
Arthropod
Arthropod
Arthropod
Arthropod
Arthropod
Arthropod
Arthropod
Arthropod
Arthropod
Arthropod
Arthropod
Arthropod
Arthropod
Arthropod
Arthropod
Arthropod
Arthropod
Arthropod
Arthropod
Arthropod
Figure S14(b)
Multiple sequence alignment of the DNA sequence in the bfr gene region for all filarial Wolbachia, with arthropod Wolbachia wCle, wOcae, wMoviF and wCfeJ sequences as references for an intact bfr gene.

### Slide 21
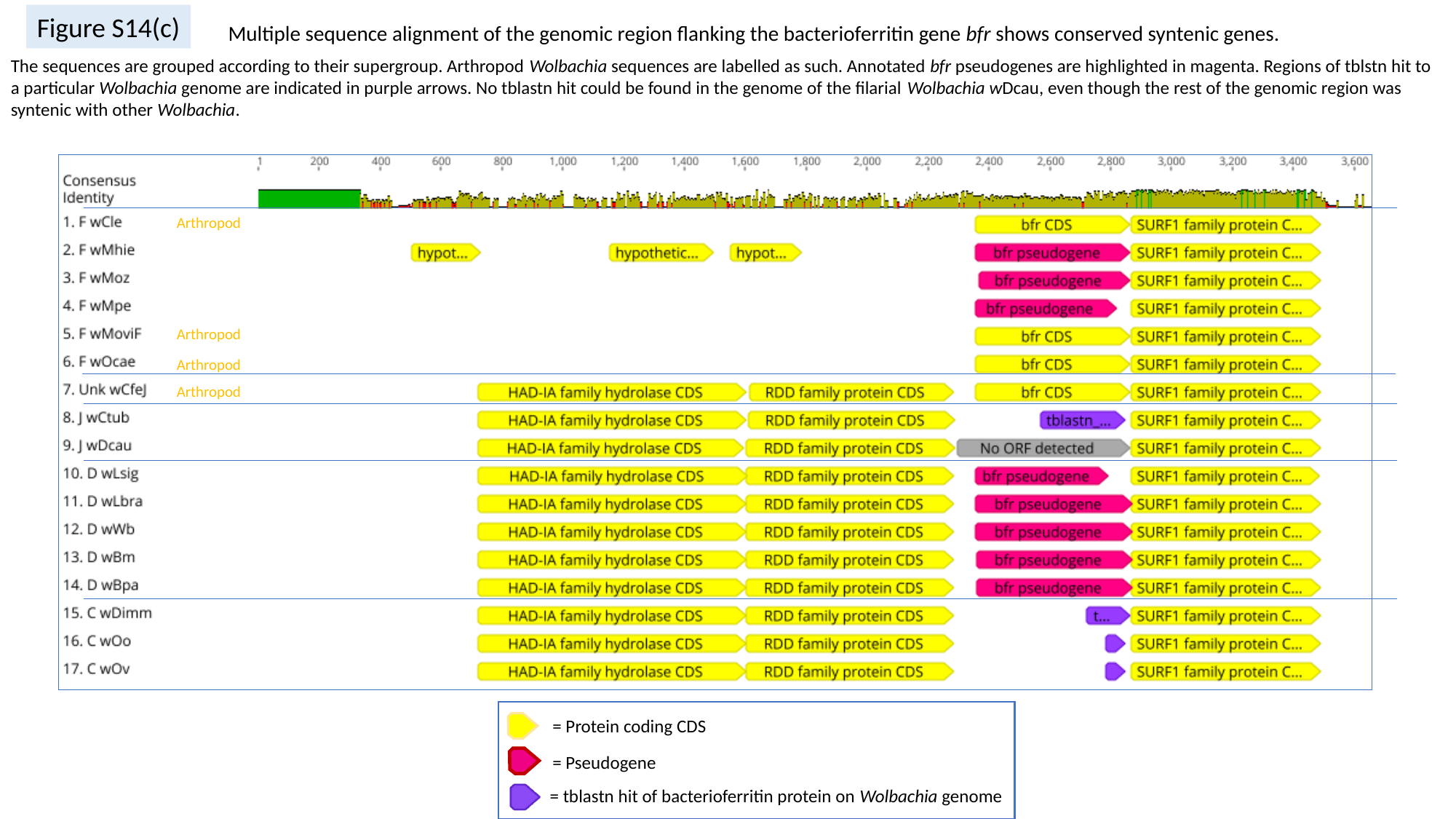

Figure S14(c)
Multiple sequence alignment of the genomic region flanking the bacterioferritin gene bfr shows conserved syntenic genes.
The sequences are grouped according to their supergroup. Arthropod Wolbachia sequences are labelled as such. Annotated bfr pseudogenes are highlighted in magenta. Regions of tblstn hit to a particular Wolbachia genome are indicated in purple arrows. No tblastn hit could be found in the genome of the filarial Wolbachia wDcau, even though the rest of the genomic region was syntenic with other Wolbachia.
Arthropod
Arthropod
Arthropod
Arthropod
= Protein coding CDS
= Pseudogene
= tblastn hit of bacterioferritin protein on Wolbachia genome

### Slide 22
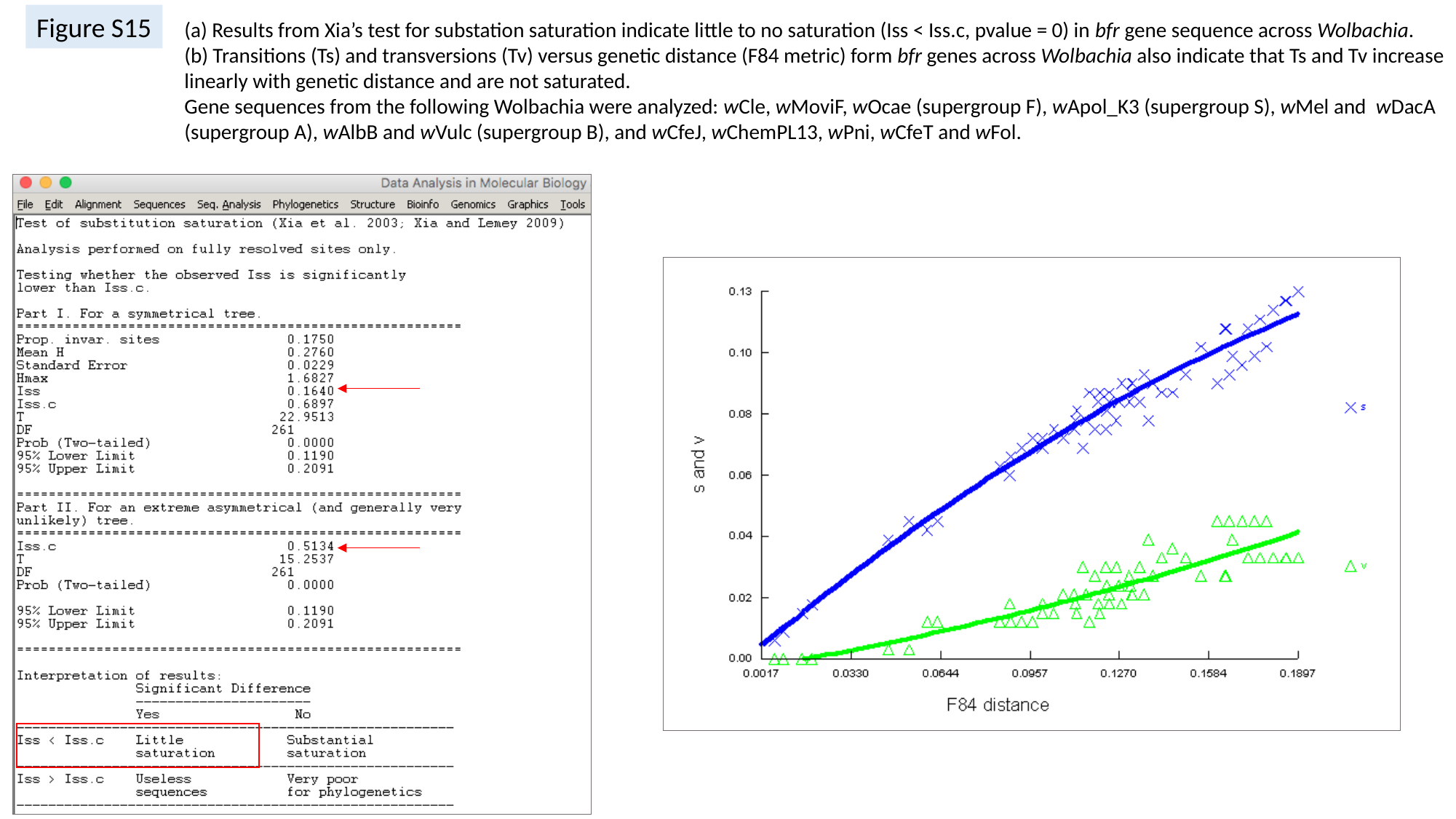

Figure S15
(a) Results from Xia’s test for substation saturation indicate little to no saturation (Iss < Iss.c, pvalue = 0) in bfr gene sequence across Wolbachia.
(b) Transitions (Ts) and transversions (Tv) versus genetic distance (F84 metric) form bfr genes across Wolbachia also indicate that Ts and Tv increase linearly with genetic distance and are not saturated.
Gene sequences from the following Wolbachia were analyzed: wCle, wMoviF, wOcae (supergroup F), wApol_K3 (supergroup S), wMel and wDacA (supergroup A), wAlbB and wVulc (supergroup B), and wCfeJ, wChemPL13, wPni, wCfeT and wFol.

### Slide 23
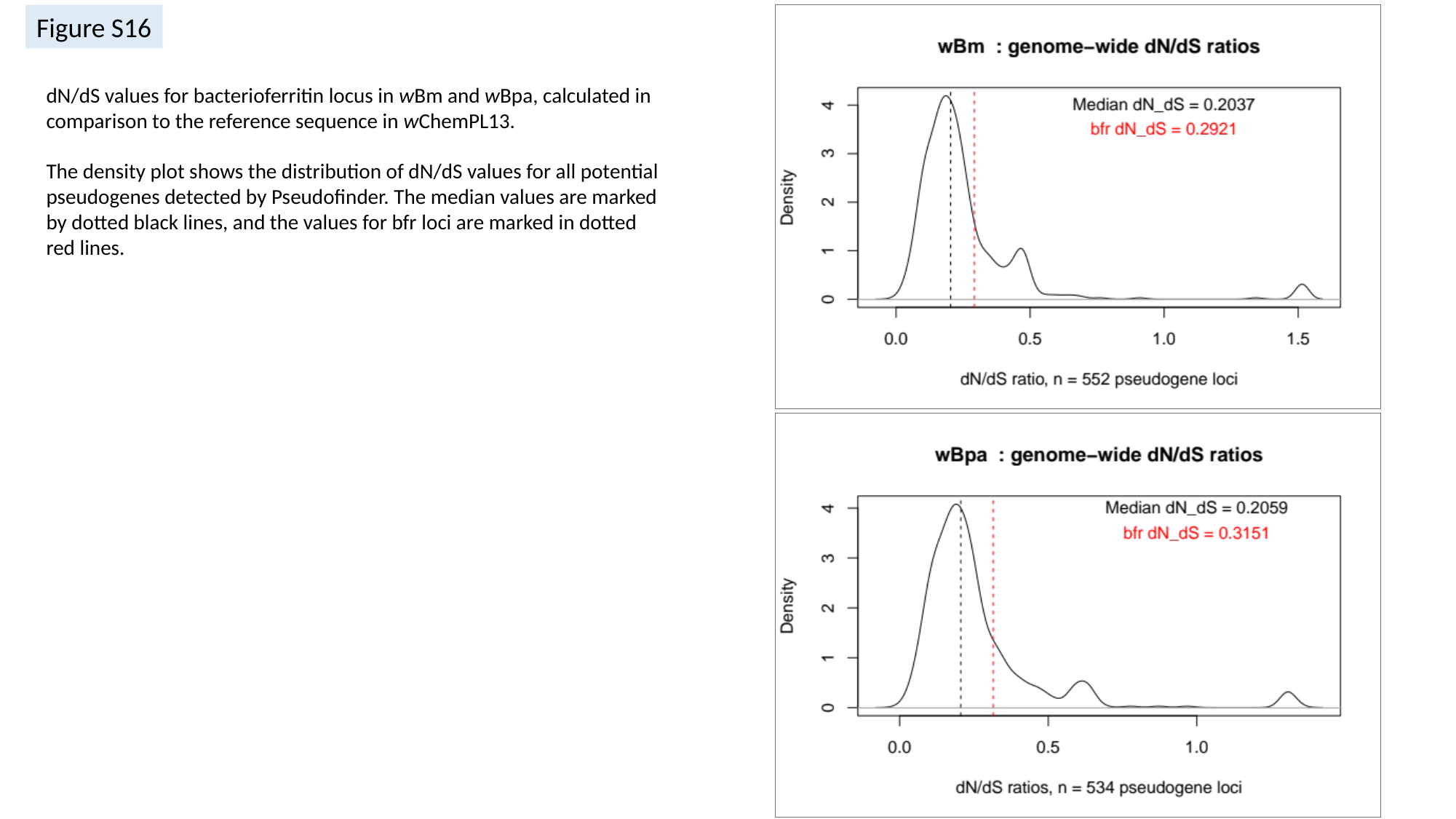

Figure S16
dN/dS values for bacterioferritin locus in wBm and wBpa, calculated in comparison to the reference sequence in wChemPL13.
The density plot shows the distribution of dN/dS values for all potential pseudogenes detected by Pseudofinder. The median values are marked by dotted black lines, and the values for bfr loci are marked in dotted red lines.
